# Development of an antibody fused with an antimicrobial peptide targeting *Pseudomonas aeruginosa:* a new approach to prevent and treat bacterial infections

**DOI:** 10.1101/2022.12.28.522163

**Authors:** Kenneth Johnson, James C. Delaney, Thomas Guillard, Fanny Reffuveille, Jennifer Varin-Simon, Kai Li, Andrew Wollacott, Eric Frapy, Surin Mong, Hamid Tissire, Karthik Viswanathan, Faycal Touti, Gregory J. Babcock, Zachary Shriver, Bradley L. Pentelute, Obadiah Plante, David Skurnik

## Abstract

The increase of emerging drug resistant Gram-negative bacterial infections is of global concern. In addition, there is growing recognition that compromising the microbiota, through the use of broad spectrum antibiotics, may affect patient health in the long term. Therefore, there is the need to develop new -cidal strategies to combat Gram-negative infections that would consider these specific issues. In this study, we report and characterize one such approach, the antibody-drug conjugates (ADCs) that combine (i) targeting a specific pathogenic organism through a monoclonal antibody with (ii) the high killing activity of antimicrobial peptides. We focused on a major pathogenic Gram-negative bacterium associated with antibacterial resistance: *Pseudomonas aeruginosa* and designed an ADC by fusing an antimicrobial peptide at the C-terminal end of the V_H_ and/or V_L_-chain of a monoclonal antibody, VSX, that targets the core of *P. aeruginosa* lipopolysaccharide (LPS). This ADC demonstrated appropriately minimal levels of toxicity to mammalian cells and rapidly kills *P. aeruginosa* strains through several mechanisms while protecting mice from *P. aeruginosa* lung infection when administered therapeutically. Furthermore, we found that the ADC was synergistic with several classes of antibiotics. This approach described in this study may result in a widely useful strategy to target specific pathogenic microorganisms without augmenting further antibiotic resistance.

**Author Summary:** The increasing of emerging drug resistant bacterial infections is a worldwide issue and infections caused by antibiotic resistant Gram-negative pathogens are particularly concerning. In addition, there is now growing recognition that disruption of the microbiota, through the use of broad spectrum antibiotics, may affect patient health in the long term. Therefore, there is the need to develop new -cidal strategies to combat Gram-negative infections while preserving the microbiota and also avoid enhancement of antibiotic resistance. We report and characterize here one such approach by using a specific monoclonal antibody associated with the potent killing activity of antimicrobial peptides in the form of an antibody-drug conjugate (ADC). The selected pathogenic bacterium was *Pseudomonas aeruginosa,* that presents numerous markers for both innate and acquired antibiotic resistance. The ADC lacked significant cytotoxicity against mammalian cells and was shown to be effective both *in vitro* and *in vivo* against *P. aeruginosa*.

## Introduction

Antimicrobial resistance is a serious, and growing, public health threat (1). The Centers for Disease Control and Prevention (CDC) estimates that in the United States, more than 2.6 million people are infected each year with antibiotic-resistant microorganisms, with at least 44,000 dying as a result (2) Of the various resistant human pathogens, Gram-negative bacteria, particularly the carbapenem-resistant *Enterobacterales* (CRE), the multi-drug resistant (MDR) *Pseudomonas aeruginosa* and *Acinetobacter baumannii* are among the most concerning. *P. aeruginosa* is intrinsically resistant to many antibiotics, limiting treatment options. Furthermore, the acquisition of resistance elements leading to MDR and even pan-resistant strains has created a public health concern with potentially untreatable *P. aeruginosa* strains (3). The CDC in its 2019 report designated MDR *P. aeruginosa* as a “Serious Threat” (2) and the World Health Organization in 2017 classified carbapenem-resistant *P. aeruginosa* as one of two “Priority 1: Critical Threats”. In addition, carbapenem-resistant *P. aeruginosa* strains were recently reported to be more fit and virulent *in vivo* (4, 5). This emerging situation warrants urgent development of new types of treatments and/or approaches to either prevent or treat *P. aeruginosa* infections (6).

Several classes of antibiotics are able to elicit rapid bactericidal effect with a greater than 99.9% reduction of the bacterial counts within four hours at peak concentrations (7). However, this very high killing ability is also associated with several shortcomings. First, these treatments induce strong selective pressure such that their use can invariably lead to the rapid emergence and dissemination of antibiotic resistance (8–10). Second, broad spectrum antibiotics act not only on the pathogenic strains, but also target the host microbiota, altering quickly and sometimes persistently its taxonomic, genomic and functional capacities, with potential negative consequences for the patient (11, 12). Thus, there is a need to develop novel targeted strategies to treat pathogenic organisms, particularly Gram-negative pathogens, with high killing abilities but with as few of these limitations as possible.

To this end, we describe the development and characterization of a new strategy to treat bacterial infections, even those caused by MDR or pan-resistant strains, combining the unique specificity of a monoclonal antibody (referred hereafter to as VSX (13)) with the direct-acting antibacterial activity of an antimicrobial peptide (AMP). While the principle of antibody-drug conjugate has recently been described to treat bacterial infection, to date, these first reports are mainly employing antibody-antibiotic conjugates (14). In this work, for the first time to our knowledge, we present an antibody conjugated with an antimicrobial peptide enabling a direct bactericidal activity against Gram negative bacteria by targeting their outer membrane. Our approach combines therefore the programmability of adding a strong antimicrobial activity with the benefits of a specific approach associated with the use of an antibody. To exemplify this approach, we focused on *P. aeruginosa* (15).

We find that our VSX-AMP constructs, henceforth referred to as antibody-drug conjugates, ADC, function with both direct bactericidal activity and effector function through the Fc domain of the antibody. We show here that such a construct demonstrates potent and selective activity *in vitro* and *in vivo*, demonstrating specific killing activity with little to no non-specific cytotoxicity to mammalian cells. Additionally, ADC constructs based on VSX and an antimicrobial peptide, employ a direct-acting effect at the outer membrane surface, which does not require internalization of the ADC, thereby circumventing the need for an agent that must pass through the double membrane of Gram-negative pathogens. Taken together, the data presented here demonstrate that our ADC constructs provide a therapeutic option for managing *P. aeruginosa* infections, promoting antibiotic stewardship and sparing the host microbiota.

## Results and Discussion

### Selection of an Antibody

VSX is an antibody that engages the inner core of LPS, including phosphorylated heptose, that is conserved across *Pseudomonas* species including *P. aeruginosa* (13). By engaging such a highly conserved site on a predominate antigen, with approximately one million copies at the outer membrane (16), VSX has the potential to be a broad spectrum immunotherapy for *P. aeruginosa* infection. As disclosed in more detail in (13), to arrive at VSX, we identified a starting mouse antibody and, using structure-guided approaches, engineered it to optimize contacts to the inner core glycan as well as improve the antibody’s drug-like characteristics. Importantly, these previous studies identified that VSX is able to bind to a wide range of *Pseudomonas* species, including *P. aeruginosa* (both rough and smooth variants), suggesting to us that it could form the basis of a construct that could target the outer membrane of Gram negative organisms.

As a first step in the process of manufacturing an VSX-AMP construct, we characterized the biological activity of the VSX antibody itself, to establish a baseline for additional studies. In particular, we examined the ability for VSX to kill *P. aeruginosa* in the presence of polymorphonuclear leukocytes (PMNs) and complement. VSX demonstrated activity in vitro using the opsonophagocyotsis killing assay (OPKA, **Figure 1A**) against the reference strain PAO1 previously used in OPKA assays (17). In vivo, we used a mouse model pneumonia that we previously reported (4). As shown **Figure 1B**, VSX injected intraperitoneally four hours post infection was able to significantly protect mice infected by *P. aeruginosa* in our acute lung infection model (P=0,04. Log-Rank test).

**Figure 1.**
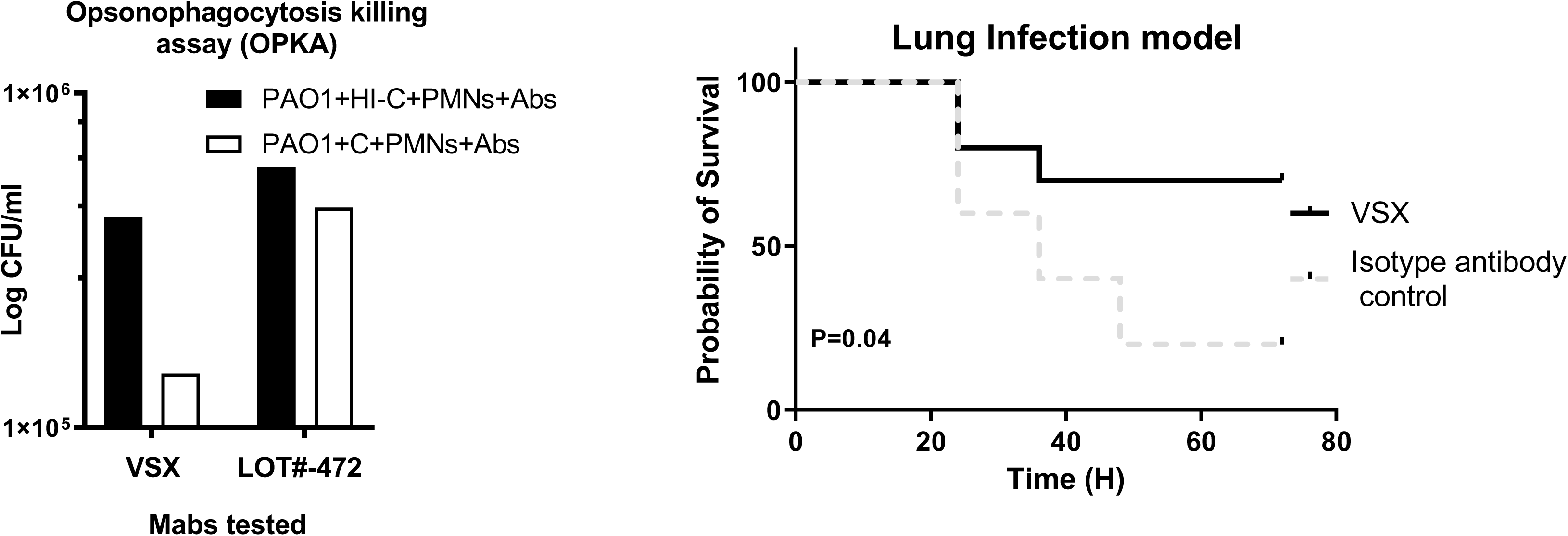
In vitro and in vivo activity of the Monoclonal antibody (Mab) VSX targeting *P. aeruginosa* core LPS. **(A)** *In vitro killing of P. aeruginosa* PA14 mediated in the presence of PMN and Complement (OPKA) by VSX. The lot#-472 is representative of a Mab able to bind *P. aeruginosa* without any detectable OPKA. C=Complement. HI C=Heat Inactivated Complement. PMN=polymorphonuclear leukocytes. Abs=Antibodies. **(B)** *Acute lung infection model*. Challenge dose: 2×10^6^ CFUs. Inoculation: intra nasal, 10^6^ CFUs in each nostril. Mice=10/group (two experiments with 5 animal/group each time). Mab were injected intraperitoneally 4 hours post infection. Dose of the Mabs: VSX=15mg/kg. Control Mab (against *Clostridioides difficile*): 15mg/kg. P. Value=0,04 measured by Log-Rank test.

### Selection of an AMP to be conjugated with VSX

To improve the in vitro and in vivo VSX killing abilities (**Figure 1**), we sought to arm VSX with direct killing activity, without the need for recruitment of either complement or PMNs (18), through the addition of a bactericidal antimicrobial peptides (AMP). We chose to focus on AMPs due to their rapid killing activity and the proven efficacy of their mechanism of action, which approximated that of the last line of defense of our current antibiotics, the polymyxins (19).

To identify an optimal AMP to be used for our antibody-drug conjugate, we completed a robust structure-activity campaign to identify AMPs that (1) are active at the cell surface and hence do not require internalization; (2) are bactericidal against *P. aeruginosa* and (3) have low hemolytic and cytotoxic activity. To this end, we utilized a bioinformatics-driven workflow coupled with experimental testing, to identify potent AMPs with a high therapeutic index. We implemented a screening strategy that clustered peptides based on their underlying physicochemical properties, followed by characterization of representative members from each cluster. Here we used YADAMP (http://www.yadamp.unisa.it/), a database of AMPs that contains over 2,500 sequences of peptides with reported antibacterial activity (**Figure 2A, top**). Then, clustering was performed by utilizing a K-means algorithm using calculated properties that have been reported to be relevant to antimicrobial activity (peptide length, predicted helicity, predicted hydropathy, percentage of select amino acids [Lys, Arg, Trp, Cys, His], and charge at pH 7, pH 5, and pH 9) (**Figure 2A, middle**). There are at least four distinct classes of antimicrobial peptides, based on their secondary structures: β-sheet, α-helix, extended, and loop (20). Since we are focused here on AMP activity in the context of covalent addition to an antibody, we eliminated the loop class, due to the requirement of disulfides to stabilize secondary structure. In addition, we eliminated peptides that require oligomerization to elicit cell killing activity. With these limitations in mind, the class of amphipathic α-helical peptides that act at the cell surface clearly represent a desirable peptide class for formation of an ADC. Furthermore, reports indicate that binding of approximately one to ten million AMPs per cell will induce cell killing (21). Thus, we reasoned that leveraging a multiple DAR ratio and using the VSX antibody to anchor the AMP in constant close proximity to the *P. aeruginosa* outer membrane creates a local high surface concentration of peptide allowing for possible membrane disruption and cell killing.

**Figure 2.**
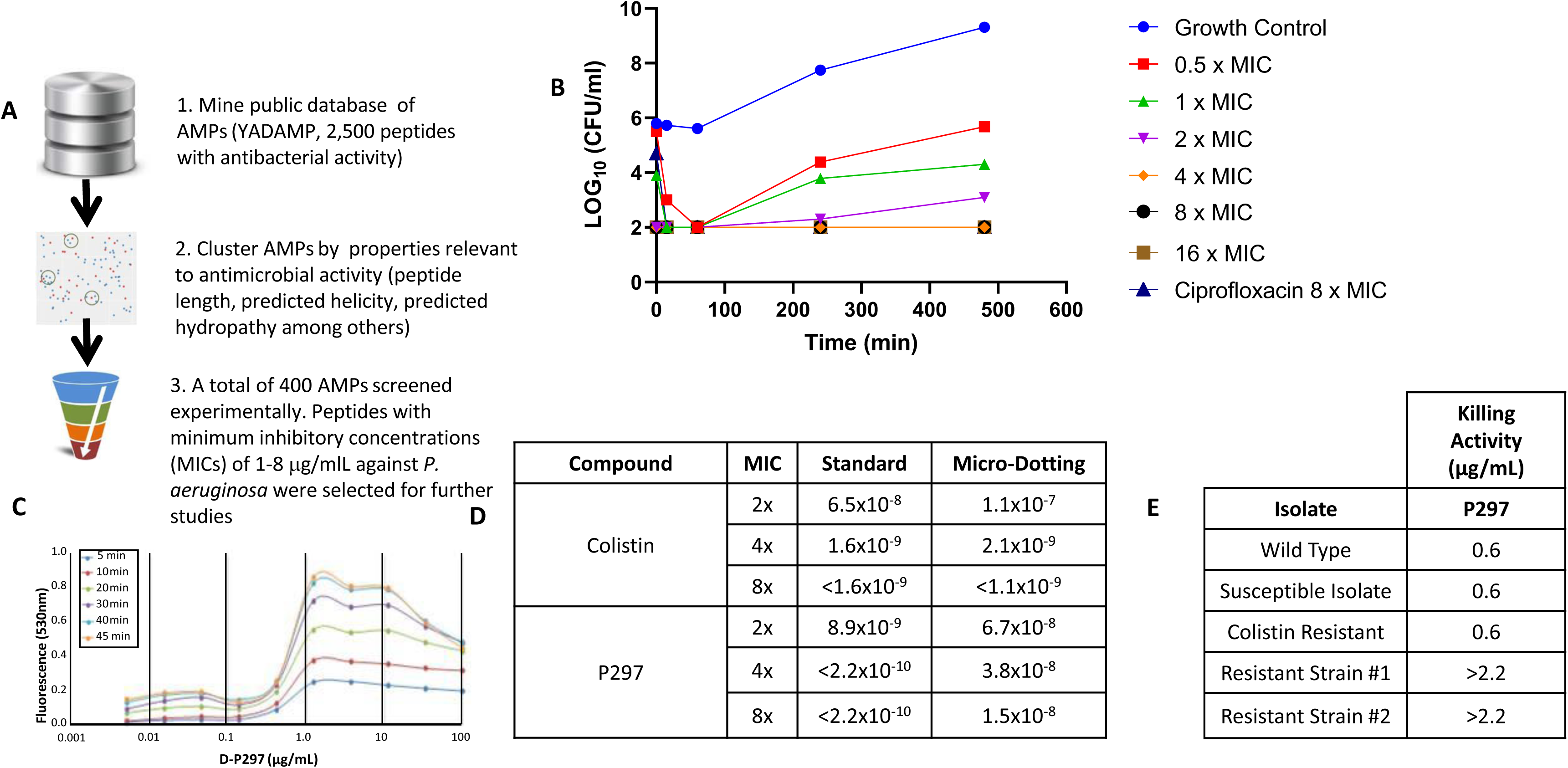
Design and screening of AMPs as potential AMPs. (**A**) Workflow for identification of an AMP to deploy in the construction of an ADC. (**B**) Antimicrobial peptide P297 showed rapid bactericidal activity in a time-kill assay using *P. aeruginosa* ATCC 27853. A growth control was compared to P297 at 0.5, 1, 2, 4, 8, or 16x MIC of peptide or to ciprofloxacin at 8x its MIC. Note that the 4x, 8x, 16x, and Ciprofloxacin 8x MICs all overlap. (**C**) *Calcein leakage*. Mechanism of action for P297 likely involves membrane disruption as assessed by measuring calcein leakage using DOPE/DOPG liposomes. Various concentrations of peptide, from 0.05– 100 µg/ml, were incubated with liposomes for a given amount of time (5-45 min). Release of calcein was assessed by measuring an increase in fluorescence at 530 nm compared to a non-peptide reference. Peptide concentrations above 1 µg/ml resulted in a measurable increase in fluorescence. (**D**) Assessment of resistance rates for P297 compared to colistin using two different procedures. (**E**) Comparison between wild type *P. aeruginosa* ATCC strain 27853 and two resistant mutants to P297 indicates differences in drug sensitivity as well as phenotypic differences. Mutant strains demonstrated decreased sensitivity to P297.

An initial set of 100 peptides was selected for experimental characterization using a panel of *in vitro* assays involving MIC testing against multiple bacterial strains, hemolytic evaluation and cytotoxicity. Once potent peptides with favorable characteristics were identified, the remaining members of the corresponding clusters were then selected for additional experimental characterization. Using this approach, we were able to dramatically reduce the number of peptides that required screening by approximately 6-fold. Over the course of the campaign, ~400 peptides were screened experimentally (**Figure 2A, bottom**).

### Identification the AMP P297

The initial screen identified several α-helical AMPs that possessed minimum inhibitory concentrations (MICs) of 1-8 μg/ml against two ATCC strains of *P. aeruginosa*: *P. aeruginosa* ATCC 27853 and *P. aeruginosa* ATCC 39324. Some AMPs also had broader activity against other bacterial species as well (**Table 1**). However, we noted that several of these initial AMPs also demonstrated lytic activity against human red blood cells. One notable exception to this trend was the α-helical sub-class of the cathelicidin family of AMPs. Cathelicidins are a family of structurally diverse AMPs that exert potent antibacterial activity and act as multifunctional effector molecules of innate immunity (22, 23). Results from multiple sequence variants from this family are outlined in **Table 1**. In particular, one member of this class, cathelicidin-BF (24), highlighted by one derivative - P297, demonstrated potent MIC values against multiple *P. aeruginosa* ATCC strains, low hemolytic activity, and a resistance to killing mammalian cells. Time-course experiments with P297 indicated that at concentrations of four times the MIC or greater, the peptide was able to rapidly reduce *P. aeruginosa* titers greater than 1000-fold, confirming that the peptide was bactericidal and not merely bacteriostatic (**Figure 2B**). Consistent with rapid onset of action, activity for P297 in a killing assay (see *Materials and Methods*) was higher than in a more traditional MIC assay (EC_50_ in the killing assay of 0.14-0.28 µg/ml compared to 2-4 µg/ml in an MIC assay).

**Table 1.** in vitro Activity and Toxicity of Representative Peptide Variants

| Sample | Sequence | MIC (µg/ml) |  |  |  | Hemolysis |  | Human Serum | Cytotox |
| --- | --- | --- | --- | --- | --- | --- | --- | --- | --- |
|  |  | *P. aeruginosa*<br>ATCC<br>27853 | *P. aeruginosa*<br>ATCC<br>39324 | *E. coli*<br>ATCC<br>25922 | *S. aureus*<br>29213 | MLC/<br>MIC | PLC/<br>MIC | hsMIC/MIC | CC<sub>50</sub><br>(µg/ml) |
| P261 | GGGGIGKFLKKAKKFGKAFVKILKK | 16 | 8 | 4 | 32 | > 8 | > 8 | 2 | 250 |
| P265 | GGGLLGDFFRKSKEKIGKEFKRIVQRIKDFLRNLVPRTES | 32 | 16 | 32 | > 128 | > 4 | 1 | > 4 | 90 |
| P267 | GGGGRFKRFRKKFKKLFKKLSPVIPLLHLG | 4 | 4 | 4 | 16 | > 32 | > 32 | 0.5 | 70 |
| P271 | GGGGLRKRLRKFRNKIKEKLKKIGQKIQGLLPKLA | 16 | 8 | 4 | 16 | > 8 | 4 | 2 | 250 |
| P292 | GGGHTASDAAAAAALTAANAAAAAASMA | > 128 | > 128 | > 128 | > 128 | ND | < 1 | ND | 250 |
| P293 | GGGGLRRLGRKIAHGIKKYGPTILRIIRIAG | 8 | 2 | 4 | 4 | 16 | 4 | 4 | 70 |
| P294 | GGGRGLRRLGRKIAHGVKKYGPTVLRRIKKYG | 64 | 2 | 4 | 4 | > 2 | > 2 | 1 | 120 |
| P295 | GGGGRFKRFRKKFKKLFKKLSPVIPLLHLG | 4 | 4 | 4 | 16 | > 32 | > 32 | 1 | 95 |
| P296 | GGGKRFKKFFKKLKNSVKKRAKKFFKKPRVIGVSIPIF | 4 | 2 | 4 | 16 | > 32 | 32 | NA | 400 |
| P297 | GGGKFFRKLKKS VKKRAKEFFKKPRVIGVSIPIF | 4<br>(1.0) | 2 (0.5) | 4 (1.0) | 32<br>(8.3) | > 32 | > 32 | 2 | 780 |
| Ctl. | Ceftazadime | 1<br>(1.8) | 1 (1.8) | 0.25<br>(0.5) | 4 (7.3) | 16 | 8 | 1 | 550 |

MIC, minimum inhibitory concentration, numbers in parentheses are molar equivalents; MLC, mean lytic concentration; PLC, partial lytic concentration; hsMIC, MIC in human serum; CC<sub>50</sub> is the concentration at which 50% cytotoxicity of mammalian cells is observed; ND, not determined; NA, no activity in human serum.

### Characterization of the AMP P297

Given its activity profile, we sought to confirm the overall structure of P297 and ensure that it aligns with the peptide’s putative mechanism of action. To this end, we first employed circular dichroism to analyze the peptide’s secondary structure. Inspection of P297’s spectrum indicated that in aqueous solution, P297 did not adopt appreciable secondary structure, with the minimum at approximately 198 nm corresponding to the π-π* transition of a random coil (**Figure S1**). However, in the presence of 40% 2,2,2-trifluoroethanol, a nonpolar solvent that has been used to promote native-like α-helical structures in peptides with intrinsic α-helix forming properties and represents, to a certain extent, the hydrophobic environment of the lipid membrane, there is a strong maximal signal at 196 nm, indicative of α-helix formation.

Additional mechanistic studies confirmed that P297 likely works through membrane disruption (**Figure 2C**, calcein leakage). In this case, model membranes were created with DOPE/DOPG liposomes and loaded with calcein, as a reporter (25). When liposomes were subjected to P297, there was rapid disruption of the lipid layer, resulting in release of dye, as measured by fluorescence. This release was concentration dependent; furthermore, time-course studies indicated that membrane disruption was rapid, reaching completion <45 minutes.

In addition to the above structure-activity studies, we sought to understand the impact of P297 administration on *P. aeruginosa*, in particular, the ability of the organism to develop resistance to it. Resistance assessment was conducted by two different methods (see *Methods*). For both methods, we compared the frequency of emergent resistance to P297 to that of polymyxin B (colistin). For P297, we noted similar or lower frequency of mutations, on the order of 10^−8^ to 10^−10^, for P297 compared to polymyxin B (**Figure 2D**). During our resistant rate determination assays with P297, a mutant strain was isolated which demonstrated significant resistance as determined in killing assays and MIC determinations (**Figure 2E**).

Mutations that confer antibiotic resistance often involve modifications of the bacteria which can lead to sub-optimal biological functions in the cell. To test if there was such a fitness cost associated with acquisition to the resistance to P297, a competitive fitness assay was run between the wild type *P. aeruginosa* ATCC 27853 and its mutant resistant strain. Briefly, we mixed 10^6^, 10^5^, and 10^4^ of each strain. At 24 hours, we plated 50 μl of serial dilutions of the mixed cultures onto blood agar plates to determine CFU/ml (see the methods section for more details). We found that the relative fitness value for the resistant strains was 0.855, comparing favorably to values reported for resistant strains to the majority of antibiotics, where a significant fitness cost was associated with a mean fitness value of 0.88 for resistant mutants (26).

Finally, we interpreted the mechanistic and resistance studies in the context of specificity, which is critical to the ultimate ADC construct. To this end, P297 exhibited a relatively high specificity, or therapeutic index, in our *in vitro* assays. One key metric that we employed, and that previously has demonstrated to be a sensitive assay of toxicity is red blood cell (RBC) hemolysis (27). For this assessment, we employed both the mean lytic concentration (MLC), that is, the concentration of peptide that induced 100% hemolysis as well as the minimal concentration at which red blood cell hemolysis is first observed (defined as partial lytic concentration, PLC). With both measurements, the levels of P297 at which RBC hemolysis was observed were 10-fold greater than the MIC. Furthermore, the cytotoxicity of P297 to the representative mammalian cell 293T was approximately 200-fold greater than the MIC.

These results, taken together, confirm that P297 is (i) an α-helical, bactericidal peptide that is able to rapidly kill *P. aeruginosa, (ii)* that it has a relatively high barrier to resistance development, (iii) that a fitness cost was associated with acquisition of the resistance to the AMP finally (iv) P297 had a therapeutic index. Therefore, P297 was selected for further peptide design.

### Improvement of P297 and selection of D297

Further analysis of P297 indicated that, while it is highly active and specific *in vitro*, in the presence of human or mouse serum the activity of P297 decreased substantially over time (**Figure S2**). LC-MS analysis of P297 in the presence of serum demonstrated that the peptide was likely adsorbing to serum components, presumably proteins, and was also being cleaved by serum proteases - its measured half-life was on the order of 20 minutes. Therefore, we set out to modify the properties of P297 to make it more amenable to long exposure times *in vivo*, especially given that antibody levels can be sustained for days. As a first modification, we replaced the L-amino acids of P297 (which adopts a right-handed helix) with D-amino acids (**D297**), a substitution which has been shown previously to decrease protease sensitivity and increase serum stability of peptides (28, 29). Circular dichroism analysis of D297 confirmed that it adopted a left-handed helix in the presence of 2,2,2-trifluoroethanol.. *In vitro* analysis of D297 indicated that it possessed equivalent potency against *P. aeruginosa* compared to P297 with the added benefit that the D-version was stable in serum (**Figure S2**). As with P297, D297 had low hemolytic and cytotoxic activity, demonstrating specificity for bacterial membranes. Thus, by converting to the D-peptide sequence we were able to retain bactericidal activity, enhance serum stability and preserve the lack of hemolytic and cytotoxicity of the AMP. Finally, we confirmed that the D297 peptide had activity against an MDR strain of *P. aeruginosa*. For this assessment, *P. aeruginosa* strain ATCC 2108 was selected; this strain has resistance to most carbapenems and cephalosporins and intermediate resistance to third generation fluoroquinolones. Despite its resistance profile, D297 demonstrated equivalent activity in the MIC assay of 4 µg/ml.

### Construction of VSX conjugates

Having selected VSX as antibody and D297 as AMP, we aimed to produce and test multiple antibody-drug conjugate (ADC) constructs of VSX-D297, using an enzymatic ligation strategy employing Sortase A (SrtA, **Figure 3A**). The SrtA method (30) involved recombinantly expressing VSX with a SrtA-ligatable tag and then enzymatically coupling the tagged VSX with the chemically synthesized D297 (30). Antibody conjugates, ADCs, were produced as either C-terminal variants of the heavy chain (HC), or light chain (LC) or both chains (dual). Conjugates with HC or LC attachment had a drug-to-antibody ratio (DAR) of ~2 (greater than 1.8 as assessed experimentally by mass spectrometry) or ~4 (greater than 3.6 as assessed experimentally by mass spectrometry) for the dual conjugate (**Figure S3**). For quality control purposes, our produced ADCs were all analyzed by mass spectrometry (LC-MS), SDS-PAGE and SE-HPLC. The couplings were efficient, routinely converting at > 90% per site as determined by LC-MS. D297 was attached in all cases with a 30 amino acid (GS)_15_-linker, to maintain flexibility, between the terminal HC/LC residue and the sortase signal sequence. As a consequence to this flexibility, SE-HPLC analysis showed a homogenous product with no signs of aggregation (**Figure 3B**). This was considered as important, as the risk of aggregation is a common concern with ADCs.

**Figure 3.**
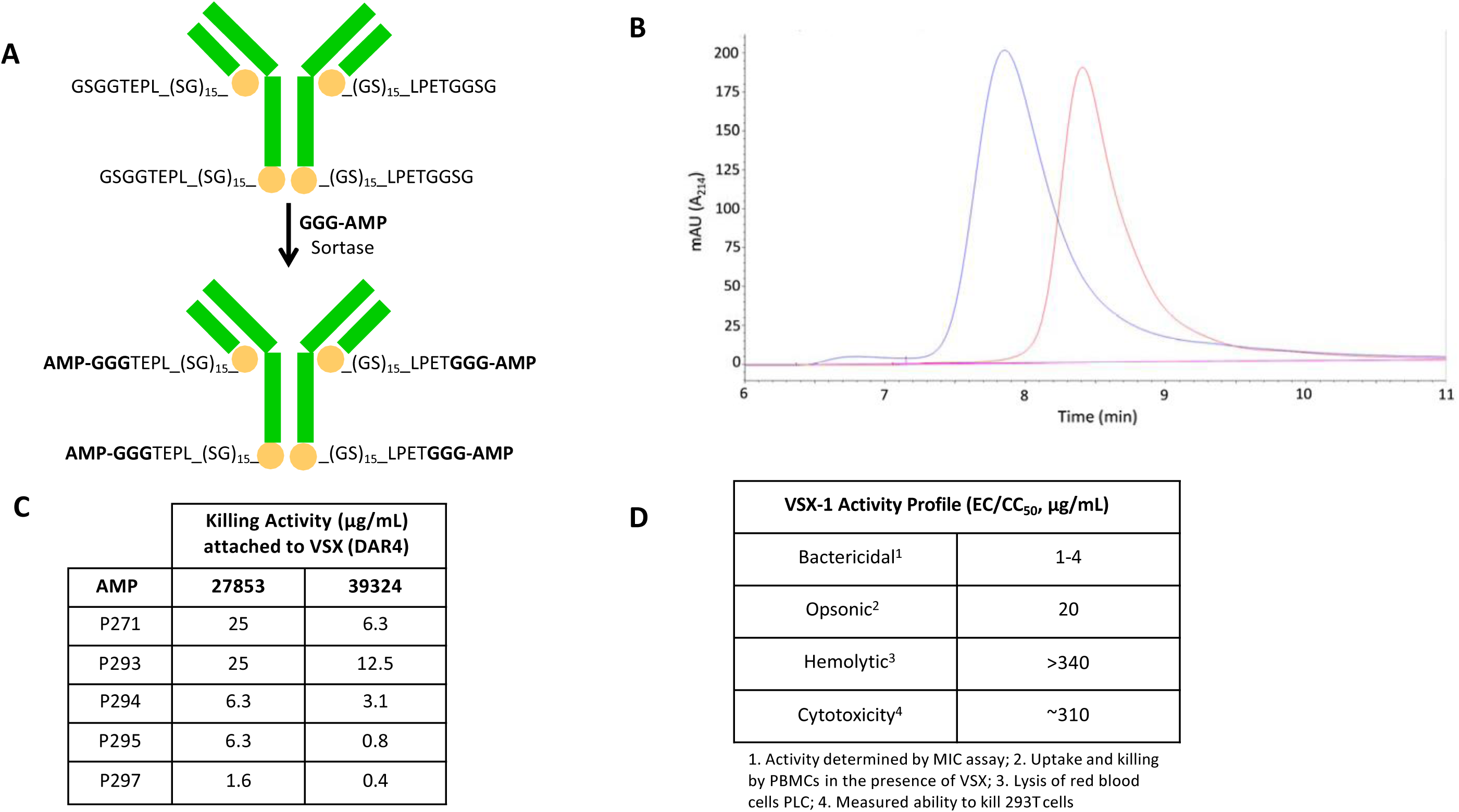
Synthesis, characterization and in vitro evaluation of VSX conjugates. (**A**) Synthesis of antibody-AMP conjugates using sortase ligation. VSX was expressed in Expi293 cells with a (GS)_15_ flexible linker and a sortase acceptor tag (LPETGGSG) present at the C-terminus of both the light chain and the heavy chain. Then, a (GGG)-modified antimicrobial peptide AMP (AMP, for example P297) was covalently added via incubation with recombinantly produced sortase for a target DAR of 4. (**B**) Size exclusion chromatography (SEC-HPLC) of VSX (red line) and VSX conjugate (blue line) indicating an earlier shift in elution time for the modified construct compared to the starting antibody. The peak width and height is similar between the constructs indicating a relative homogeneity in sortase modification. (**C**) *In vitro* killing activity of VSX conjugates. *P. aeruginosa* ATCC strains 27853 or 39324 were treated with VSX conjugates with a DAR of ~4 containing AMP peptides 271, 293, 294, 295, or 297 and conjugate IgG concentrations that result in 50% killing are recorded. (**D**) VSX-1 (VSX with peptide P297 and a DAR of ~4) was characterized by several assays including *in vitro* bactericidal activity, opsonic activity, hemolytic activity and cytotoxic activity.

### In vitro killing and therapeutic index of the ADC VSX-1

Using the VSX antibody, a series of ADC constructs were evaluated *in vitro* for direct bactericidal activity against *P. aeruginosa*. From these initial constructs, we were able to conclude the relative order of activity for the sites of conjugation to be (HC + LC) > HC > LC. The most potent ADC, hereafter referred to as VSX-1, contained the VSX antibody and a DAR of four with AMP D297 ligated to the C-terminus of each HC and LC (HC+LC). VSX-1 exhibited potent bactericidal activity in a killing assay against two ATCC strains of *P. aeruginosa* (**Figure 3C**). Consistent with the results obtained with D297, VSX-1 had a high therapeutic index with minimal hemolytic activity and negligible cytotoxic activity against 293T cells (**Figure 3D**). To confirm the maintenance of each component within the ADC, the activity of the antibody, the peptide, and VSX-1 was assessed in a mix culture of *P. aeruginosa*, *E. coli* and *K. pneumoniae* (**Figure S4**). Consistent with its activity profile the P297 AMP alone (**Figure S4, left panel**) showed no specific killing with killing detected for both *P. aeruginosa* and *E. coli.* Consistent with its requirement of the presence of complement or PMNs, the antibody alone exhibited no killing (**Figure S4, panel in the middle**). In contrast to both the AMP and the antibody, the ADC specifically killed *P. aeruginosa* (**Figure S4, right panel**).

### Other in vitro properties of VSX-1

(i) *The OPKA (opsonophagocytic killing assay)*. The killing properties of the ADC were also evaluated in an OPKA to confirm intactness of the antibody. The ADC was highly active at 10 μg/ml, resulting in a 10-fold bacterial reduction in the presence of heat-inactivated complement and greater than 1000-fold killing activity with non-heat inactivated complement, in the presence of polymorphonuclear neutrophils. As with OPKA assessment of VSX, the strain *P. aeruginosa* PAO1 was used (17). **(ii) *Synergy with antibiotics*.** Given that VSX-1 acts at the outer membrane of *P. aeruginosa*, we reasoned that it may demonstrate synergy with existing antibiotics, which must cross the double membrane to target intracellular targets. Indeed, we find that VSX-1 appears to increase the potency of classical anti-*Pseudomonas aeruginosa* antibiotics (*e.g.* carbapenems and polymyxin B) with a 10-fold reduction of the MIC or more (**Figure 4**). Indeed, we found that an increasing amounts of VSX-1 from 0.015 to 4 μg/mL lowered the observed MIC for meropenem from 0.5 to <0.1 μg/ml and that of colistin from ~1 μg/ml to <0.1 μg/ml (**Figure 4**). This result is likely caused by the AMP increasing the outer membrane permeability. (**iii**) ***LPS-mediated TLR4 activation inhibition***. Finally, other additional mechanisms of protection that could be mediated by VSX were also explored. Specifically, it has been reported that shedding of LPS may cause pathophysiological manifestations upon hyperstimulation of the host immune system – initiated by activation of toll-like receptors (TLRs) which, in turn, can lead to septic shock (31). Given this pathophysiology, an anti-LPS antibody could play a vital role in neutralizing LPS by limiting LPS shedding, promoting its serum clearance and inhibiting the LPS-mediated binding to TLR and concomitant activation of the immune system. Consistent with this pathway as well as the binding site of VSX, a HEK-blue reporter assay that measures the activation of TLR4 receptor by free LPS demonstrates that VSX inhibits LPS-mediated TLR4 activation (**Figure S5**).

**Figure 4.**
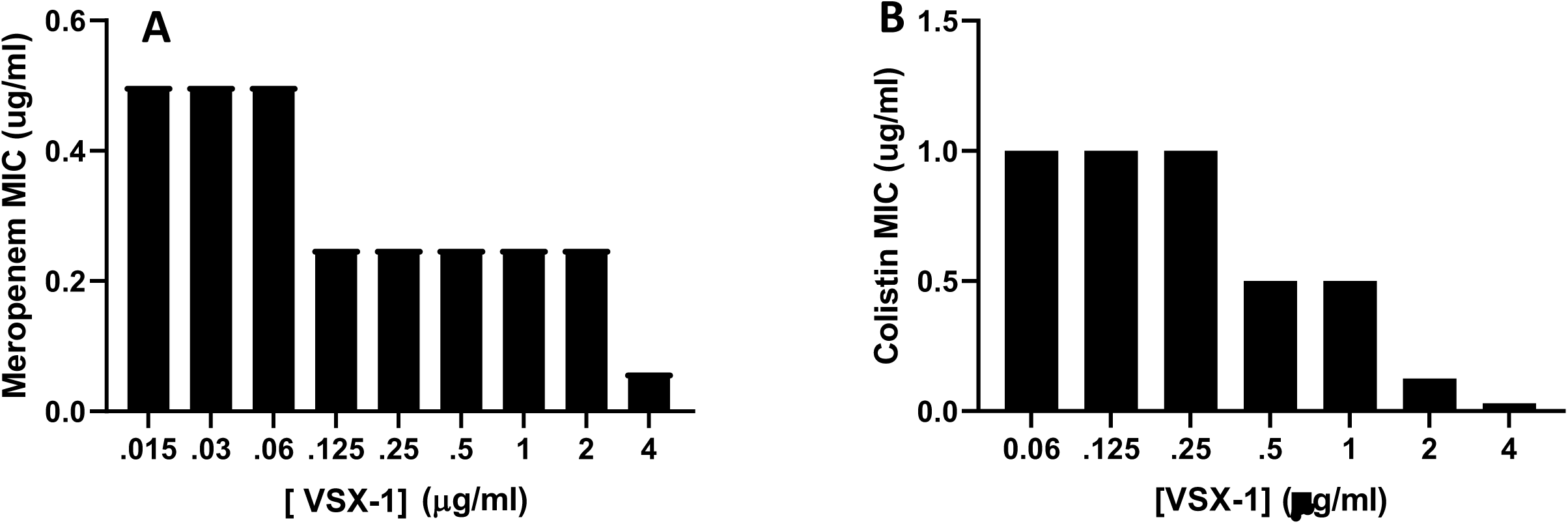
Synergy of P297 with different antibiotic classes. MIC for both meropenem and colistin was determined alone and then varying concentrations of P297 were titrated to measure the effect on MIC towards P. aeruginosa ATCC 27853. (**A**) Examination of MIC for meropenem in the presence of different concentrations of P297. (**B**) Same as (**A**), except colistin was used as the antibiotic. Not all antibiotics demonstrated synergy; for example, the aminoglycoside tobramycin did not exhibit an enhanced MIC in the presence of P297.

### In vivo activity of VSX-1 in animal models of infections caused by P. aeruginosa strains

To investigate the *in vivo* activity of VSX-1, we focused on lung infection models for several reasons. First, the lung is a common site for *P. aeruginosa* infection, leading to community/hospital-acquired pneumonia, which is associated with problematic outcomes, with high mortality rates, up to 30-60% in some studies (32). As such, it presents one area of likely clinical use for a pathogen-specific ADC. Second, given the compartmentalization of the lung from the bloodstream, it also represents a high bar to demonstrate efficacy for a systemically administered antimicrobial.

Consistent with the *in vitro* data, VSX-1 demonstrated *in vivo* efficacy in multiple animal models of *P. aeruginosa* infection. A murine neutropenic lung infection model using *P. aeruginosa* (ATCC 27853) was previously described (33) and was initially employed to explore the proof of concept that our ADC VSX-1, an antibody based therapeutic, would work in vivo even in an immunocompromised host because it is conjugated to another antimicrobial agent. Infected animals were treated intranasally with VSX-1 construct resulting in statistically significant reduction in the bacterial load in the lung both when co-administered (**Figure 5A**) and when dosed therapeutically (**Figure 5B**). Notably, antibody alone showed no bacterial reduction, supporting the selection of an ADC with direct bactericidal activity as a preferred mechanism, particularly in an immunocompromised setting. Building on this initial data, VSX-1 was evaluated in a murine immunocompetent model (**Figure 5C**) using the same approach that we previously reported and that we also used in this study when testing the antibody VSX alone (**Figure 1B**): the acute lung infection model with the *P. aeruginosa* strain PA14 (4). While VSX alone was able to significantly protect the infected mice, the survival of infected animals treated three hours post-infection with a single intraperitoneal dose of VSX-1 at 15 mg/kg (**Figure 5C**) was more pronounced (P=0,04 and P=0,007 respectively).

**Figure 5.**
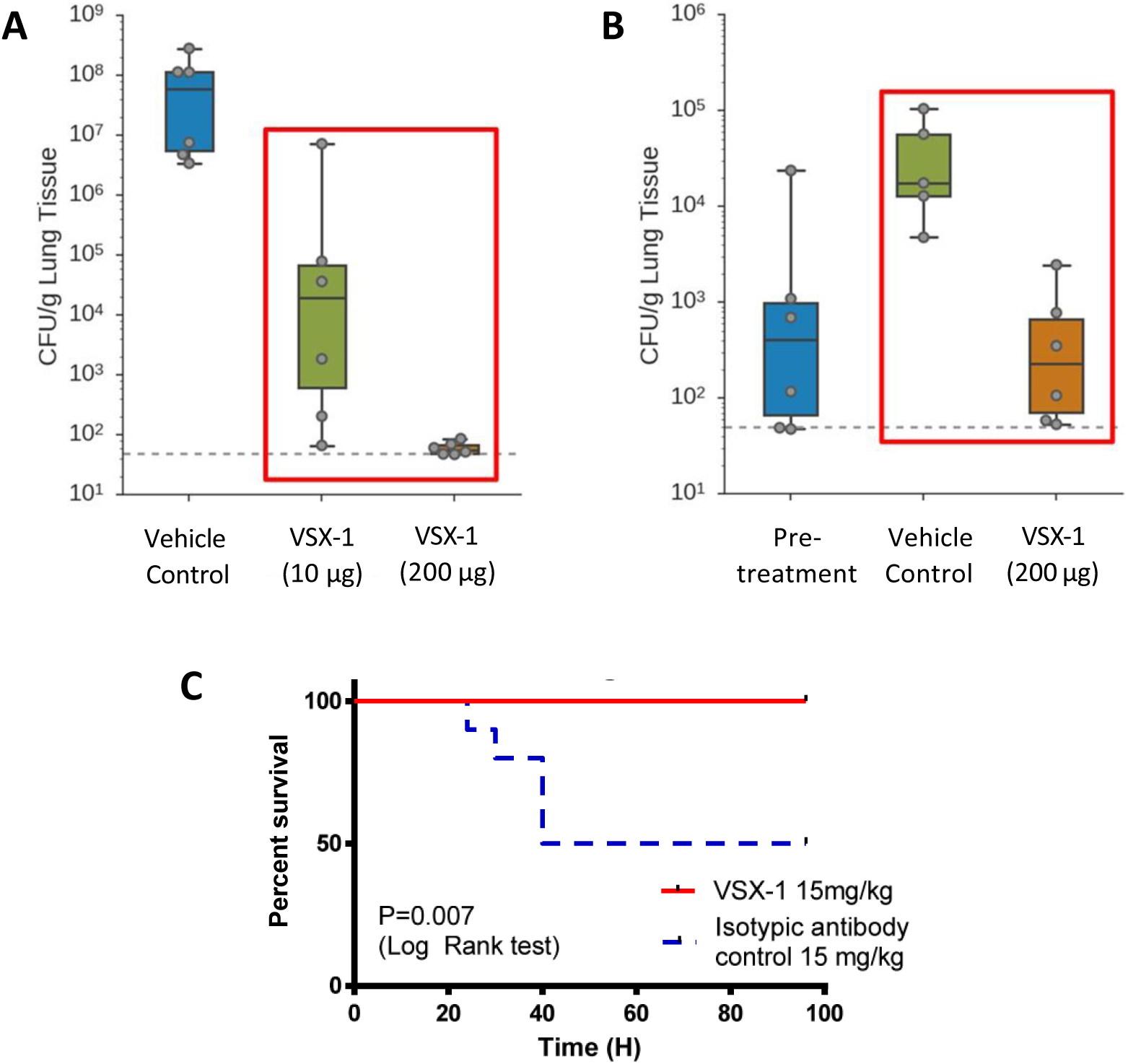
Evaluation of conjugates in in vivo models of P. aeruginosa lung infection. (**A**) Neutropenic animals co-administered VSX-1 and bacteria (ATCC 27853). CFU burden as measured in the lung at eight hours. Co-administration of 10 µg of VSX-1 resulted in a multi-log reduction in bacterial burden, with reduction to the limit of detection upon administering 200 µg of ADC. (**B**) Neutropenic animals were infected with *P. aeruginosa* (ATCC 27853) and were treated 1 hour post-infection with either vehicle or VSX-1 (200 µg). CFU burden was measured in the lungs just before treatment (pre-treatment) and in the lungs at eight hours post-injection. (**C**) Acute lung infection model with *P. aeruginosa* PA14 (2×10^6^ CFU/animal, 10^6^ in each nostril), C57/Bl6, 10 animals/group (two experiments with 5 animal/group each time), intranasal inoculation, intraperitoneal dosing (15 mg/kg) 4 hours post-infection.

The immunocompetent model used for this experiment is aggressive, with ≥ 50% of control animals succumbing to infection within 24 hours of inoculation. To confirm the contribution of the antibody component of VSX-1, a sham-conjugate was prepared using a non-*P. aeruginosa* targeting antibody (actoxumab, an human monoclonal antibody against *Clostridioides difficile* (*34*)). The actoxumab-D297 conjugate showed no survival benefit in the immunocompetent lung infection model, validating the selection of a *P. aeruginosa* surface-targeting antibody. In all, the protection observed with the VSX-1 conjugate was highly encouraging and warranted the further exploration of this ADC as a potential treatment for *P. aeruginosa* lung infection.

### Optimization of the ADC: VSX-2

The promising first generation VSX-1 construct was further evaluated *in vivo*. Despite complete protection in *P. aeruginosa* infection models, we noted that VSX-1 displayed somewhat compromised biodistribution in mice relative to the parent antibody (**Figure 6A**). Based on previous studies with ADCs, the AMP properties and DAR have been shown to lead to compromised bioavailability *in vivo* with similar constructs (35). Indeed, we find that a DAR of two with the AMP D297, while not as potent, demonstrates improved bioavailability in the mouse with increased circulating levels after 1h, 24h and 72 hours post-administration, compared to VSX-1 (**Figure 6B**). This observation carried though to other compartments as well, with increased bioavailability in the lung after 1h (**Figure 6C** top left panel for VSX-DAR2, **Figure 6A** middle top panel for VSX-DAR4).

**Figure 6.**
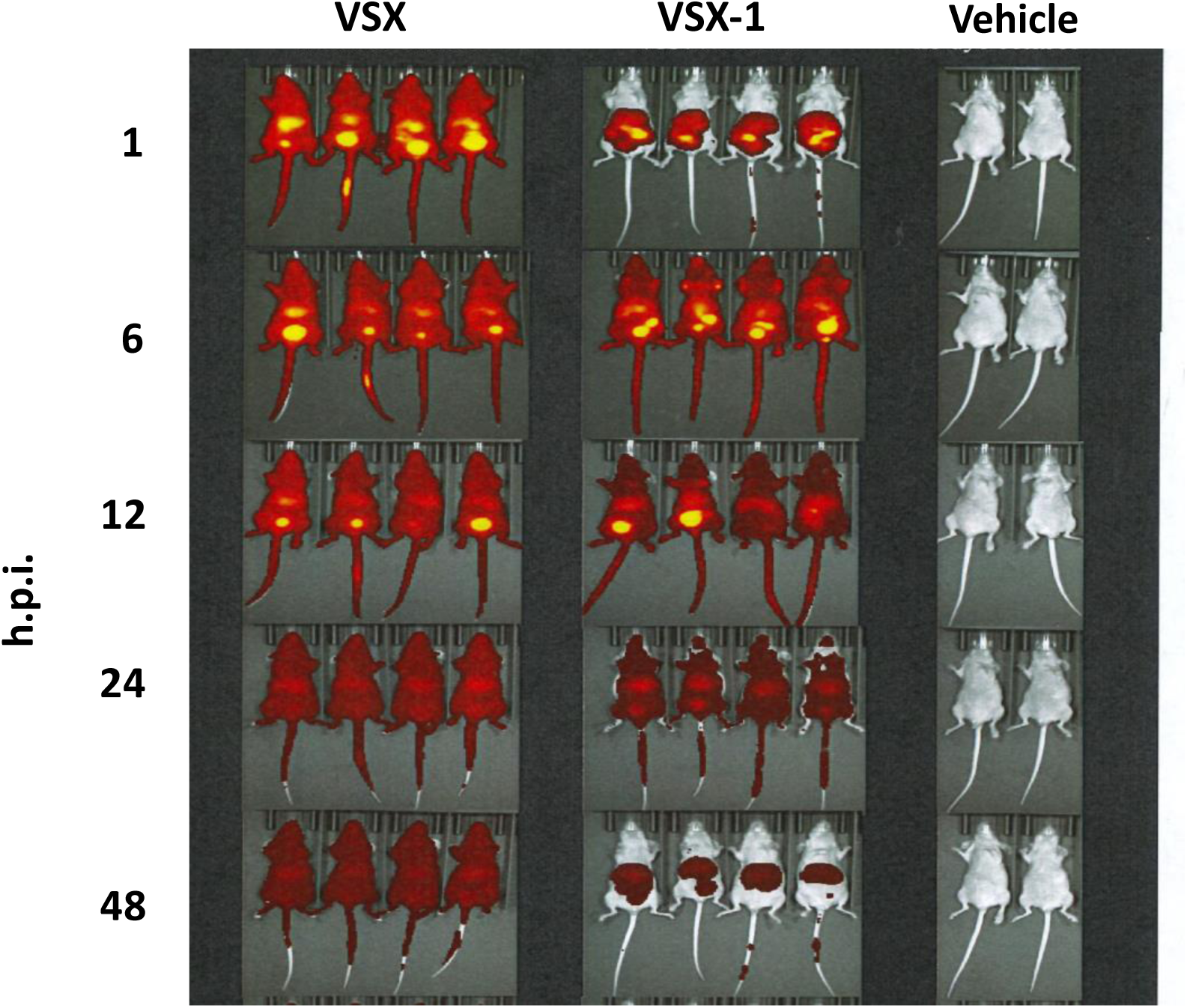
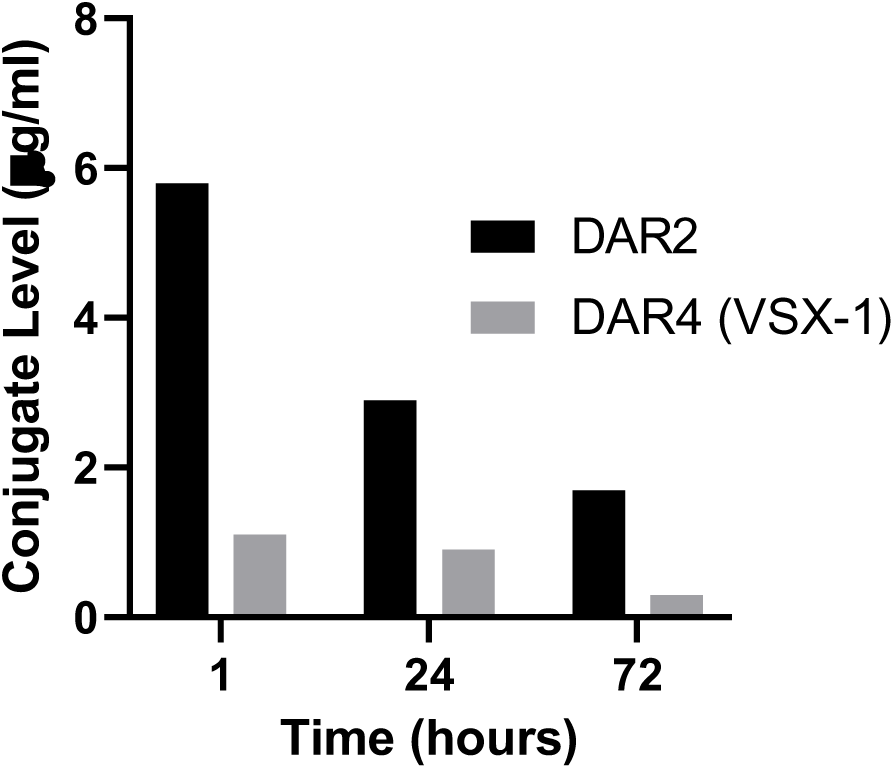
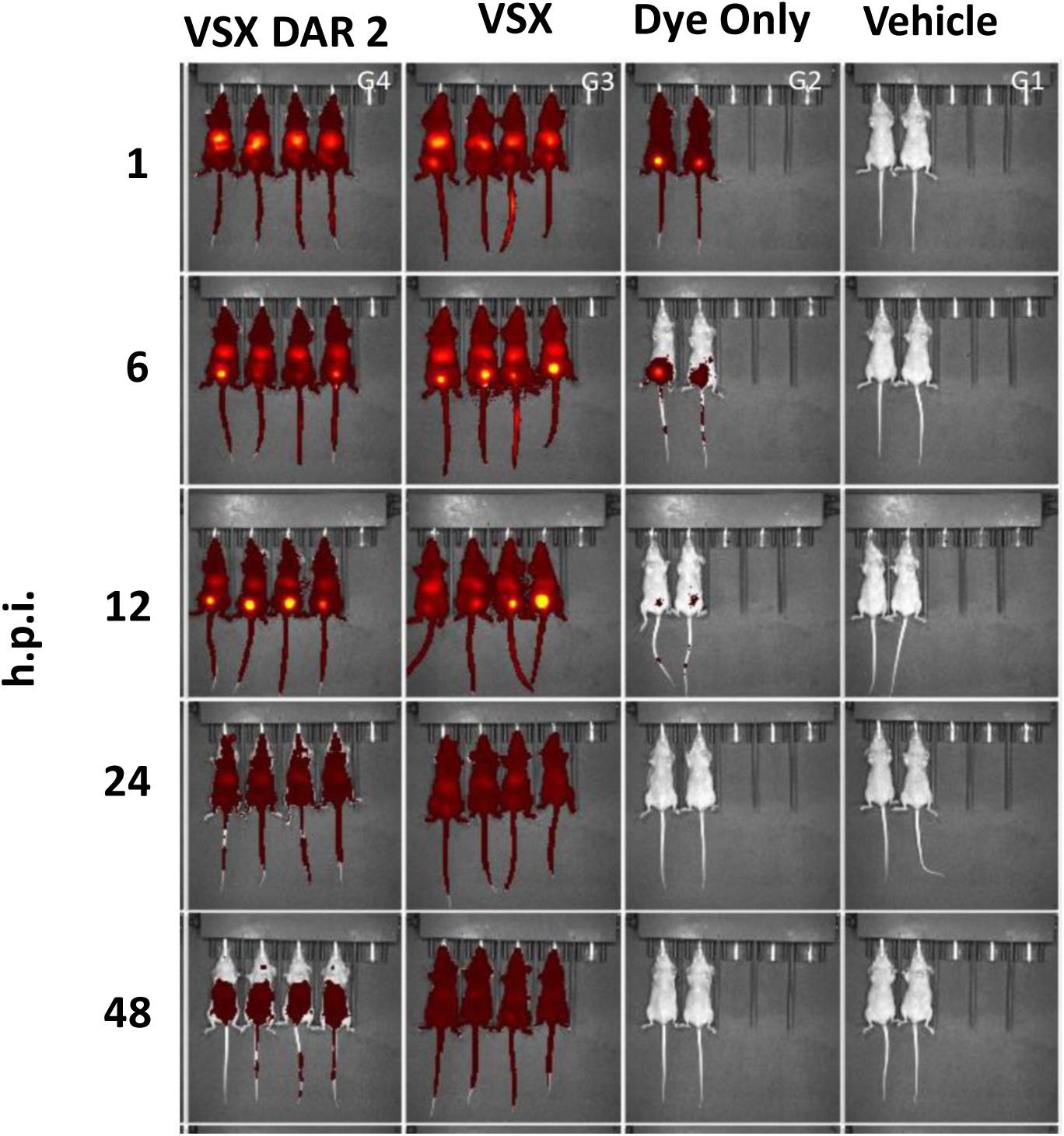
Biodistribution VSX-1. (**A**) *In vivo* imaging of VSX-1 biodistribution. Labeled VSX (antibody alone) or VSX-1 (ADC) were administered to animals IP and followed by imaging out to 48 hours post-injection (h.p.i.). Both agents distributed through the animal, to all perfused organs, but VSX-1 had a faster elimination, with little agent evident at 48 h.p.i. (**B**) Analysis of serum levels by ELISA of VSX-1 (DAR4) (grey bars)at 1, 24, and 72 hours post-injectionand DAR2 constructs (black bars). (**C**) Same imaging procedure as (**A**) but with a DAR 2 demonstrates better biodistribution and half-life of the ADC.

Therefore, to optimize the biodistribution properties of VSX conjugates as a potential therapeutic, we carried out a focused set of studies on AMP charge, AMP hydrophobicity and DAR. Starting from the P297 sequence, we screened a set of charge variants and variants with globally reduced hydrophobicity. Unexpectedly, L-P369 (GGGKLLRKLKKSVKKRAKELLKKPRVIGVSIPL), containing 5 phenylalanine to leucine substitutions, emerged as a more potent peptide and, when conjugated to the VSX antibody, retained bactericidal activity and demonstrated little cytotoxicity to mammalian cells (**Figure 7A**). As with D297, P369 (containing either D- or L-amino acids) demonstrated potent MIC activity across several *P. aeruginosa* strains, including the MDR ATCC strain 2108 (MIC = 8 µg/ml). Indeed, VSX-2, containing a DAR of 2 with AMP D-369 ligated to the C-terminus of each HC, also demonstrated protection in the *P. aeruginosa* lung infection model (**Figure 7B**). Thus, it appears that we can modify both the activity and the PK characteristics of the ADC through modification of the peptide characteristics and the DAR.

**Figure 7.**
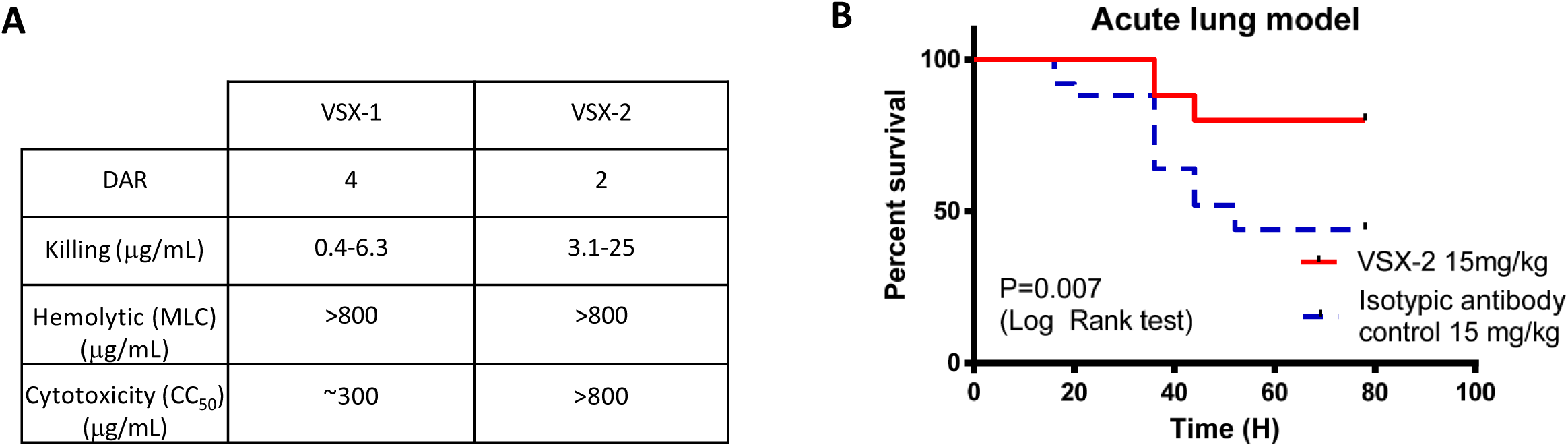
In vitro and in vivo assessment of VSX-2. (**A**) Activity of VSX-2 compared to VSX-1 as measured by the (bacterial) Killing Assay, hemolysis of red blood cells (mean lytic concentration (MLC)) and toxicity to mammalian cells (CC_50_). The bactericidal activity of VSX-2 is similar, but slightly lower, than that of VSX-1, with a lower DAR, and with a similar inability to lyse RBCs or kill mammalian cells. (**B**) VSX-2 has similar activity *in vivo* in the acute lung infection model with *P. aeruginosa* PA14 (2 × 10^6^ CFUs/animal, 10^6^ CFUs in each nostril), C57/Bl6, 25 animals/group (five experiments with 5 animal/group each time), intranasal inoculation, intraperitoneal dosing (15 mg/kg) 4 hours post-infection.

Finally, *P. aeruginosa* is also responsible of chronic infections, for instance in Cystic Fibrosis patients (lung infections) or in wound infections for example. In these chronic infections, the treatment is often complicated by the production of biofilm by *P. aeruginosa*. Therefore, as a last step we also tested the activity of our ADC VSX-2 against biofilm grown *P. aeruginosa* both in an inhibition/prevention setup, but to also against mature biofilms (eradication/treatment setup, **Figure 8**). We used a dynamic model with continuous and very low flow of minimal medium with *P. aeruginosa* biofilms grown for 48 hours, at 37°C in flow chambers. While VSX-2 applied 24h after formation of a dynamic biofilm was able to totally eradicate *P. aeruginosa*, the efficacy of the treatment decreased overtime (**Figure 8A-B**). Prevention of biofilm by our ADC (injection of VSX-2 in the flow cell system during bacterial inoculation) was highly successful in our model (**Figure 8 C-D**)

**Figure 8:**
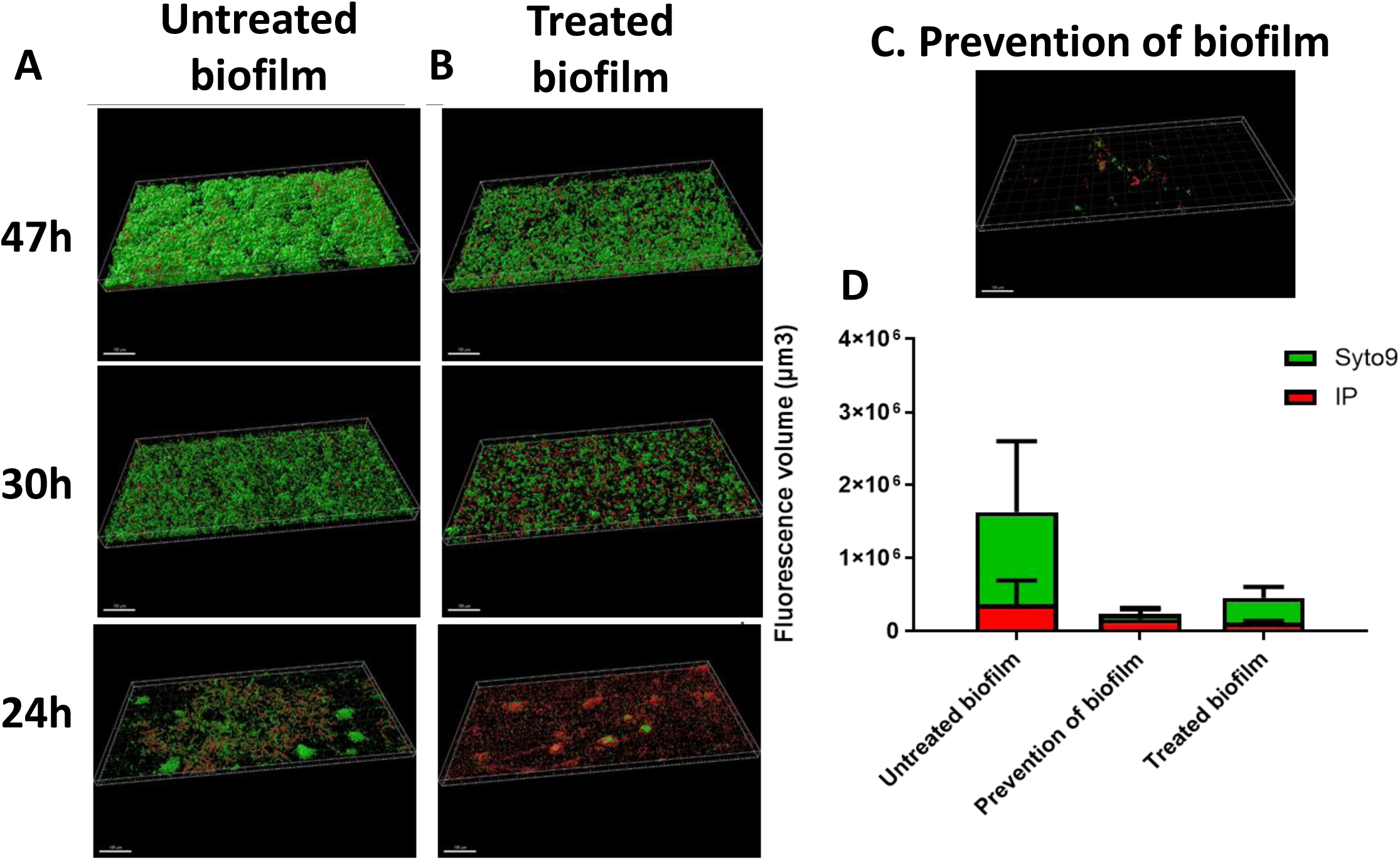
Prevention and treatment of *P. aeruginosa* PA14 biofilm in dynamic model marked with Syto9 (green) and PI (red) fluorochromes. (**A-B**). **A**: untreated biofilm (control) and **B**: Treated biofilm with 6 µg/mL of DAR2 (VSX-2) during 1h at 47h (top), 30h (middle) and 24h (bottom). (**C**) Prevention of biofilm, injection of 6 µg/mL of DAR2 (VSX-2) during bacterial inoculation at time 0. (**D**) Fluorescence volume of biofilm biomass with repartition of Syto9 and PI staining; scale = 100 µm. Acquisition of images by fluorescent microscopy of two joined fields of one sample and calculation by Imaris software. Experiments were repeated three times.

The results of this study taken together, report a new approach to treat bacterial infections, including infections caused by multi-drug resistant strains through the use of an Antibody-Drug conjugate, ADC, which can act at the outer membrane of Gram negative organisms, specifically *P. aeruginosa*. Our selected antibody targets a conserved glycan structure on the surface of *Pseudomonas aeruginosa* (13). It has been engineered in an ADC format to have direct bactericidal activity. Notably, unlike ADCs that have been constructed for oncology indications (36) and recently for Gram-positive organisms (37), internalization and release of the AMP is not required for activity: we present here construction of an ADC that acts exclusively at the outer membrane surface. The resulting ADC demonstrates significant *in vitro* activity against a variety of strains of *P. aeruginosa* and has demonstrated *in vivo* efficacy in an aggressive lung infection model.

While these data demonstrate initial proof-of-concept that an ADC approach can work for a Gram negative organism, there are some limitations to this study. First, while preliminary work has indicated that the frequency of bacterial resistance to the peptides and ADCs is low (~10^−8-10^, roughly equivalent with colistin), additional work will be required to further characterize such resistant mutants. Second, while the biodistribution of the VSX-1 ADC is sufficient to exhibit *in vivo* activity and is significantly improved with the redesigned VSX-2, neither construct exhibited biodistribution or PK that is comparable to the antibody alone. Finally, additional studies and other ADC constructs are required to identify whether other Gram negative organisms, beyond *Pseudomonas* can be targeted with such an approach.

Overall, with our ADC strategy, we augment the bactericidal activity of specific antibodies. By bringing together two optimized components, a specific human anti-*P. aeruginosa* antibody and an optimized antimicrobial peptide, the resulting ADC has therapeutic properties superior to either component alone. The data and approach presented here offer an alternative strategy for the development of antimicrobials that complements existing and ongoing efforts in small molecules and biologics (38).

## Materials and Methods

### P. aeruginosa strains

All strains used in the study and their origin are listed table S1. *VSX Antibody.* The fully humanized antibody, VSX, has been identified to target the inner core of *P. aeruginosa* LPS, specifically the phospho-diheptose. A complete description of this antibody is presented in Elli, *et. al.* (13).

### Sortase Tagging

A sortase A recognition sequence (LPETGGSG) was placed at the C-termini of either the heavy chain (DAR_theoretical_ = 2) or both the heavy and light chains (DAR_theoretical_ = 4) of the VSX monoclonal antibody. Prior to ligation, the VSX antibody was buffer exchanged from 1x PBS into sortase buffer (150 mM NaCl/50 mM Tris, pH 7.5) using 30 kDa spin diafiltration units (Amicon Ultra 15). Sortase A was from BPS Bioscience (San Diego, CA). VSX was ligated with the transpeptidase enzyme and a GGG-sortase donor peptide (added to the reaction as an aqueous solutions at 10 or 20 mg/ml), thereby replacing the GGSG sequence on the antibody with full length peptide. Ligations were performed in sortase buffer using 20 mol of peptide per mol VSX antibody (which was at 1.5 mg/ml), 10 mM CaCl_2_, 5.8 µg/ml Sortase A. Samples were kept in the dark at room temperature for 18 hours, followed by quenching via dilution to 10 ml total volume in PBS and immediate purification by Protein A FPLC. Conjugation efficiency was determined by Q-TOF mass spectrometry using a reduced antibody prepared by heating 5 µg of sample at 65 °C for 15 min in 10 mM DTT.

### SEC-HPLC

VSX and VSX conjugate were dissolved at 1 mg/ml and 10 ul were injected onto a BioSep SEC-s3000 Phenomenex column (300 × 7.8 mm) run @ 1 ml/min in 0.1M NaH_2_PO_4_ buffer at pH 3 on an Agilent 1100 series system with 214 nm UV monitoring.

### Assessment of Peptide AMP Stability in Serum

Normal human serum (NHS) (Sigma S-7023) was thawed, diluted in water, centrifuged at 13,600 × g for 10 minutes and the supernatant was warmed to 37 °C in a water bath. Twenty microliters of each test article were placed in a 2.0 ml round bottom microfuge tube. Two milliliters of diluted NHS were added to each tube which was immediately vortexed, followed by transfer of 200 µl to a fresh microfuge tube placed at 37 °C in a rotating rack. Samples were harvested and processed at various time-points up to 6 hours by quenching with 40 ul of 15% trichloroacetic acid (TCA), chilling on ice for 15 minutes, centrifuging at 13,600 × g for 10 minutes and storing the supernatant at −20 °C until analysis.

### Assessment of Antimicrobial Activity

MICs and a Killing Assay were performed in this study.

MICs were determined according to CLSI guidelines, using 2-fold serial compound dilutions, in 96-well microtiter plates. Briefly, compounds were diluted in water across a mother plate then 2 µl was stamped to assay plates, one plate for each strain to be tested. Bacterial strains were sub-cultured overnight on agar plates at 37 °C. Overnight plates were used to prepare 0.5 McFarland cultures in 0.85% saline. These concentrated cultures were diluted 1:200 in growth media to approximately 5 × 10^5^ cells/ml. All assay plates received 100 µL diluted culture per well. All plates were placed at 37 °C overnight. After 18 hours, the plates were assessed using a mirrored plate reader and reflected incandescent light. The MIC is defined as the lowest concentration of compound that inhibits growth by at least 80%. Wells at and above the MIC should appear void of growth when visualized.

To assess microbial killing (*killing assay*), bacterial cells were grown aerobically overnight on agar plates at 37 °C. Overnight plates were used to seed 30 ml cultures of growth media in 250 ml vented flasks. Cultures were grown aerobically at 37 °C, shaking at 150 rpm. Growth was monitored at A_600_ and bacterial cells were harvested at mid-log growth. Ten milliliters of culture were pelleted at 4000 xg for 10 minutes and washed one time with PBS + 1% BSA (PBSA) before re-suspending in 2 ml PBSA. The concentrated culture was used to seed 6 ml tubes of PBSA to an OD giving a concentration of 1 × 10^8^ cells/ml. Cultures were diluted to 1×10^4^ cells/ml in PBSA. Test articles were diluted in PBSA and 50 µL per concentration tested was loaded into a 96 well polypropylene microtiter plate. Fifty microliters of diluted culture were added to all test wells and no compound control wells. Plates were shaken and then incubated at 37 °C, 90 minutes, under static conditions. Ten microliters from each assay well was plated onto agar plates and incubated at 37 °C overnight. Percent killing was determined by the CFU for test wells compared to the CFU for no compound control wells. The EC_50_ reported is the lowest concentration of a compound which causes > 50% colony reduction compared to the no compound control for the strain being tested.

### RBC Hemolysis

Test articles are diluted 2-fold in water across a 96 well polypropylene mother plate leaving one no compound control. Two µl from each well is stamped onto a 96 well polystyrene assay plate. A red blood cell suspension is made by mixing 0.2% defibrinated sheep blood (Hardy Diagnostics DSB100) in PBS. All assay plates received 100 µl/well of the red blood cell suspension. Plates are then incubated at 37 °C overnight. The titer for RBC hemolysis was defined as the lowest concentration of compound that completely prevented the formation of a red blood cell pellet perceptible by eye.

### Mammalian Cell Cytotoxicity

On day one, cells from an established 293T human cell line are seeded onto 96 well flat bottom white plates with 10,000 293T cells/well. On day two, dilution plates are prepared by diluting compound in cell growth media to two times the final starting concentration and then carrying out serial doubling dilutions across the plate. Media from the cell growth assay plates is then aspirated and 50 µl from the compound dilution plates is transferred to the assay plates. 50 µl of fresh media is then added to all wells of the assay plate. Plates are incubated at 37 °C, 5% CO_2_ for three days. Then, CellTiter Glo (Promega G7570) is reconstituted and mixed 50:50 with growth media. Media from the assay plates is aspirated and 100 µl of the CellTiter Glo/growth media mix is added to all wells. After five minutes, luminescence is read and % inhibition *vs.* compound concentration is plotted. The CC_50_ is defined as the cytotoxic concentration reducing viable cell number by 50% compared to cells in media lacking the test article.

### LPS Neutralization

A cell-based LPS neutralization assay was developed and optimized. HEK-Blue LPS detection Kit2 (Invivogen) was used to investigate the ability of VSX to neutralize endotoxin activity of extracted *P. aeruginosa* LPS in HEK-Blue cells. Endotoxin, if present in the media or standard are sensed by TLR4 leading to the activation of NF-κB and the concomitant production of SEAP in the supernatant. When supernatant is combined with QUANTI-Blue, which contains a SEAP chromogenic substrate, a purple/blue color appears and can be quantified by measuring the absorbance at 620-655 nm and extrapolating against a standard curve. Endotoxin units (EU) of 0.5 EU/ml were used to define the LPS concentration added alone or pre-mixed with VSX (0.185 - 10 µM) and activation of the NF-κB was assessed. The monoclonal antibody CDA1 does not target bacteria and was used as a control in the neutralization assays

### Mixed Microbial Assay

Bacterial strains were grown overnight on agar plates at 37 °C. Overnight plates were used to establish 0.5 McFarland Cultures in 6 ml PBS (approximately 1 × 10^8^ cells/ml). Concentrated cultures were diluted to 1 × 10^4^ cells/ml in PBS. Ten microliters of each of the diluted cultures were plated onto blood agar plates (BAPs) to determine initial concentration, check for purity, and establish strain morphology. One milliliter of each diluted culture was combined (for a total of three) and the volume was brought to 10 ml with PBS (1 × 10^3^ cells/ml of each strain). Twenty-five microliters of the mixed culture was spread on a BAP to establish the CFUs/ml for each strain at t=0. Test articles were serially diluted 4-fold in PBS with a final volume of 200 µl in 2 ml round bottom Eppendorf tubes. A no compound control was included. Two hundred microliters of mixed bacterial culture was added to all assay tubes. Tubes were vortexed and immediately 50 µl from each tube was spread on separate BAPs. Assay tubes were incubated at 37 °C, rotating between timepoints. The plating procedure was repeated at 1 hour intervals for two hours and all plates were incubated at 37 °C overnight. The following day all plates were counted, noting the CFU for each strain, distinguished by different morphology, and results were plotted as the percentage killed compared to a no compound control.

### Resistance Assessment by MIC

Resistance assessment to D297 was assessed using two, complementary methods. The first, standard protocol involves plating a high-density culture on selective plates that are at various multiples of the MIC for D297 and, as a reference, colistin. Overnight cultures were brought to an OD of 3.0 at an absorbance of 600nm (10^9^ - 10^10^ CFU/ml). 100ul of concentrated culture was plated in quadruplicate onto selective plates containing compound at 2, 4, and 8 × MIC. Additionally, 100ul of 10^−7^, 10^−8^, and 10^−9^ serial dilutions of the concentrated cultures were plated in quadruplicate on non-selective plates to calculate CFU/ml. All plates were incubated at 37C and counted at 24 and 48 hours. Resistance rates were calculated. Colonies growing at 2 × MIC and higher were re-plated on selective plates to confirm resistance.

Secondly, resistance to peptide D297 was determined using the Pranting protocol’s micro-dotting procedure (39). Solid phase Minimal Inhibitory Concentrations (MICs) for test articles on Tryptic Soy Agarose plates were determined by plating 100 µl of a bacterial culture with > 1 × 10^9^ CFU/ml onto selective plates containing test compound at concentrations equal to various multiples of a broth MIC previously determined using CLSI standards. The concentration of the plate leading to an 80% reduction of CFU/ml compared to a no compound control plate defined the solid phase MIC. Selective Tryptic Soy Agarose plates were prepared at concentrations of 0, 2, 4, and 8 times the solid phase MIC for D297 and, as a reference, colistin. Bacterial test strains were grown overnight, rotating at 37° C in 40 separate 10 ml Pyrex screw-capped tubes containing 3 mls Meuller Hinton Broth II, cation adjusted (MHB). Overnight cultures were spun down and re-suspended in Tryptic Soy Broth (TSB) to a density of approximately 4.8 × 10^9^ CFU/ml as determined by densitometer. Serial dilutions of representative cultures were plated to confirm cell concentrations. All cultures were plated on selective and non-selective plates by dropping 5 µl of culture onto agarose plates and letting the drops absorb. Plates were incubated at 37° C overnight. The mutation rate was calculated using the P_o_ method, −[ln(P_o_/P_tot_]/N, where P_o_ is the number of cultures (spots)with no mutants, P_tot_ is the number of cultures, and N is the number of bacteria applied in each spot..

### Generation of Resistant Mutants

Plates generated during resistance assessment using both the standard and Pranting protocols (above) were used to isolate mutant strains of *P. aeruginosa* 27853 with elevated MICs to peptide D297. One colony from the standard protocol and 5 colonies from the Pranting protocol grew at 4 × the agarose MIC for wild type *P. aeruginosa* 27853. These six colonies were picked and mixed with PBS. 5 ul drops containing approximately 2-4E+08 cells were re-plated onto D297 selective plates (1 - 4 × MIC), colistin selective plates (0.5 - 4 × MIC), and non-selective plates to determine if the mutations confer true resistance and if any cross-resistance to Colistin has been created. Plates were incubated at 37C overnight. Of the six potential mutants, 4 confirmed growth at 2 × the MIC and 1 confirmed growth at 4 × MIC on D297 selective plates. Only one of the 6 isolates grew at 2 × MIC on Colistin selective plates. Wild type control strains had no growth at 2 × MIC for either D297 or Colistin.

### Competitive Fitness Assay

A competitive fitness assay was run with the wild type *P. aeruginosa* 27853 and the resistant mutant strain. Both isolates were grown overnight on agar plates, followed by growing both plated strains independently for 24 hours in 35 mls MHB in flasks. Overnight cultures were used to make 0.5 McFarland cultures using a densitometer. 50 ul of each 0.5 McFarland culture was plated onto blood agar plates. Cultures were then mixed so that 3, 30 ml cultures in flasks would have approximately 1E+06 of each cell type (1:100 dilution), 1E+05 of each cell type (1:1,000 dilution), and 1E+04 of each cell type (1:10,000 dilution). At 24 hours, we plated 50 ul of serial dilutions of McFarland cultures and mixed cultures onto blood agar plates to determine CFU/ml. All plates were at 37C overnight. The following day, we counted all plates for CFU/ml then re-plated the mixed bacterial flasks by diluting each mixed culture to 0.5 McFarland, diluting to 1E+05 cells/ml and plating 50 ul of the diluted cultures onto blood agar plates. All plates at 37C overnight then read for CFU/ml.

To determine whether there was a phenotypic difference between wild type *P. aeruginosa* strain ATCC 27853 and its resistant mutant, both strains were plated on eight different types of selective and non-selective agar mediums. On one type of plate (TSA with 5% sheep blood) there was a discernible difference in the colony morphology between the wild type and mutant strains, with the later exhibiting a rougher colony morphology, enabling the use of these blood agar plates for the determination of the relative fitness of the two strains.

To assess relative fitness, we used the method of Lenski and colleagues (40) to estimate the selection coefficient on a genotype, *s*, from competition data (where relative fitness is given by 1 + *s*). The growth parameter for a strain is the number of doublings that it experiences over a given period of time. As such, the selection coefficient on the focal strain is defined as follows:

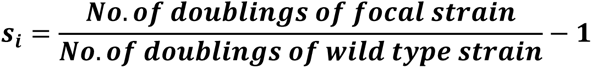

Note that *s_i_* is a unitless parameter.

### Assessment of Serum Stability

Normal human serum (NHS) (Sigma S-7023) was thawed, diluted in water, centrifuged at 13,600 × g for 10 minutes and the supernatant was warmed to 37C in a water bath. Twenty microliters of each test article at 10 mg/ml was placed in a 2.0 mL round bottom microfuge tube. Two milliliters of NHS (25% in dH_2_O) was added to each tube and immediately the tubes were vortexed and 200 ul was transferred to a fresh microfuge tube with 40 ul of 1% Trichloroacetic acid (TCA). Assay tubes were placed at 37C in a rotating rack. Additional samples were similarly harvested and processed at t = 30, 60, 120, 240, and 360 minutes. TCA tubes were placed on ice for 15 minutes and then centrifuged at 13,600 × g for 10 minutes. Supernatant from each tube was collected and frozen at −20C for analysis. Mass spec analysis was used to quantify the percent of test articles remaining intact.

### Opsonophagocytic killing assay

Opsonophagocytic Assays were performed in 2.0 ml Eppendorf tubes using a total volume of 400 ul. Four components were added in rapid succession in equal 100 ul volumes: test articles, bacterial culture, PMNs, and Human complement. Test articles were diluted in GVB +Ca +Mg (Boston Bioproducts #IBB-300X) and kept on ice until ready for use. Overnight bacterial cultures were grown in Columbia Broth + 2% NaCl (CSB), 37C, rotating at 250 rpm. Cultures were diluted to an OD at 650nm = 0.4 in GVB. Cultures underwent a second dilution, 1:200 in GVB, for a final assay concentration of 1.5E+07 cfu/ml. Human PMN were isolated with EasySep (StemCell Cat. # 19666) as per protocol from peripheral blood. PMN were resuspended in GVB to 1E+07/ml (1E+06 PMN/tube). For no PMN control tubes, GVB was used. Complement used was 20% MN8-absorbed Human C’ (single source) in GVB. Complement underwent further absorption using 200 ul C’ with 800 ul of a bacterial suspension in GVB with an OD at 650nm = 1.0. Absorption performed at 4C, 30 minutes. Cells were spun out and the process repeated with the supernatant used to resuspend a cell pellet from 800 ul cell culture. Cells were spun out and the final supernatant was filtered using 0.22um spin filters. Final complement source was 2x absorbed, 20% human complement. A small aliquot was heat inactivated at 56C, 30 minutes for a no complement control. Components, as described, were combined and 25 ul from each assay tube was removed for a t = 0 CFU determination. The assay tubes were capped and placed at 37C for 90 minutes with end over end rotation. Removed samples were serially diluted 1:10 in TSB/tween and 10ul of the 1:10 and 1:100 dilutions were plated on TSB blood agar plates allowing for sample to run down a vertical plate almost to the edge. After the 90 minute incubation the plating procedure was repeated. All plates were incubated overnight at 37C and CFU/ml calculations were performed for t=0 and t=90 minutes.

### Neutropenic Mouse Model

Animal experiments were performed in accordance with the Institutional Animal Care and Use Committee. CD1 Mice were supplied by Charles River (Margate, UK) and were specific pathogen free. Male mice were 11-15 g on receipt and were allowed to acclimatize for at least 7 days. Mice were housed in sterilized individual ventilated cages exposing the mice at all times to HEPA filtered sterile air. Mice were rendered neutropenic by immunosuppression with cyclophosphamide at 200 mg/kg four days before infection and 150 mg/kg one day before infection by intraperitoneal (IP) injection. The immunosuppression regime leads to neutropenia starting 24 hours post administration of the first injection which continues throughout the study. *P. aeruginosa* strain ATCC 27853 was used to assess *in vivo* protection. For infection, animals were first anesthetized with a ketamine/xylazine anesthetic cocktail (90 mg/kg ketamine & 9 mg/kg xylazine) via IP injection delivered at ~15 ml/kg. Anesthetized mice were infected with 0.04 ml inoculum by intranasal instillation into mouse nostrils (20 µL per nostril, 5-minute interval between each nostril administration) and were kept in an upright position on a string rack for ~10 minutes post-infection. The inoculum concentration was 2.83 × 10^5^ CFU/ml (~1.1 × 10^4^ CFU total inoculum). VSX-1 or controls were administered either intranasally (IN) or IP. The clinical condition of the animals was monitored and animals that succumbed to the disease were euthanized. The study was terminated ~24.5 hours post-infection when most of the vehicle mice were displaying significant clinical symptoms, after which the clinical condition of all remaining animals was assessed. After being euthanized by pentobarbitone overdose, mice weights were determined before the lungs were removed and weighed. Lung samples were homogenized in ice cold sterile 1x PBS using a Precellys bead beater; the homogenates were quantitatively cultured onto *P. aeruginosa* selective agar and incubated at 37 °C for 16-24 hours before colonies were counted. Data were analyzed using StatsDirect software (version 2.7.8). The non-parametric Kruskal-Wallis test was used to test all pairwise comparisons (Conover-Inman) for tissue burden data.

### Acute Lung Infection Model

The acute lung infection model was performed as previously described (4) with some minor modifications. Briefly, after general anesthesia (IP injection using ketamine and xylazine) of C57/BL6 female 6-8 week old mice, 10 μL (1×10^6^ CFU) of the reference strain *P. aeruginosa* PA14 was inoculated in each nostril to induce an acute lung infection. The inoculum was prepared from a dilution of an overnight culture of PA14 grown in Luria Bertani Broth. Mice were monitored over four days and were euthanized when they showed imminent signs of mortality including ruffled fur, lethargy, shaking, high respiratory rate, inability to move when touched or inability to right itself after being placed on its side. For the protection assays, the ADC or the control compound were injected IP at 15 mg/kg three hours after the bacterial challenge. For each experiment there was 5 animal per group. Experiments were repeated two times with VSX and VSX-1 and five times with VSX-2.

### In vivo Imaging

CR female NCI Ath/nu mice were placed on Special Diet “RD D10012Mi” for 7 days prior to study start and for the duration of the study. Animals were randomized into treatment groups based on Day 1 bodyweight with an age at Start Date of 8 to 12 weeks. VSX was conjugated with IR800 dye (LiCor, Lincoln, NE), then used in sortase reactions to make VSX-1. A vehicle control (0.9% saline) and a dye alone control group were also employed. A 5 mg/kg dose was used, and materials were injected by the IV route. Whole body imaging (dorsal and ventral) was collected at 1, 6, 12, 24, 48, 100 hours post-IV dose.

### Flow Cells/Biofilm experiments

*P. aeruginosa* biofilm were grown in a dynamic model with continuous and very low flow of minimal medium (MM) [62 mM potassium phosphate buffer, pH 7.0, 7 mM (NH_4_)_2_SO_4_, 2 mM MgSO_4_, 10 mM FeSO_4_) containing 0.4% glucose and 0.1% casamino acids].

In this dynamic model, *P. aeruginosa* biofilms were grown for 48 hours, at 37°C in flow chambers (IBI scientific, UK). The system was sterilized by pumping a 0.2 % of hypochlorite solution for 1 hour using a peristaltic pump. After a sterile water rinsing, MM was introduced for 1 hour with a rate of 20 mL/h for system stabilization. Bacteria at a concentration of 5.10^8^ CFU/mL were injected in flow cell chambers, which were flipped upside-down, without flow for 2 hours. The pump was turned on to authorize a constant rate of 0.2 mL/h of MM during 48 hours. For prevention approach, bacteria were inoculated with the VSX-2 (6 µg/mL). For treatment experiment, we injected VSX-2 (6 µg/mL) for 1 hour at three different times (24h, 30h and 47h). After 48 hours, Live/Dead BacLight bacterial viability kit (Molecular probes) at a ratio 1:5 of Syto-9 to propidium iodide is injected. Then stained biofilms were observed using a LMS 710 NLO, confocal laser scanning microscope (Zeiss). Homogeneity of the samples was checked by traversing the observation field and the most representative area was chosen for the acquisition of the image. 3D reconstructions and fluorescents volumes were generated using Imaris software. The experiment was repeated 3 times.

### Data Availability

The sequence of the VSX antibody has been deposited in GenBank. The sequences of the anti-microbial peptides P297 and P369 have also been deposited.

## Acknowledgements

We thank Dr. Gerald Pier for access to bacterial strains and advice throughout the course of this research. This work was supported by CARB-X award number: 1 IDSEP160030-01-02.

## Appendices. Supplemental Information is included (Figures S1-S6 and Table S1)

**Table S1:** Strains used in the study

| Assay | Origin | name |
| --- | --- | --- |
| MIC and Killing assays | ATCC strains | *P. aeruginosa* 27853, 39324, 2108. *E. coli* 25922, *S. aureus* 29213, *K. pneumoniae* 33495 |
| OPKA | Infection and Immunity (17) | *P. aeruginosa* PA01 |
| Acute Lung Model | Science Translational Medicine (4) | *P. aeruginosa* PA14 |
| Neutropenic Mice | Antimicrobial Agents Chemotherapy (33) | *P. aeruginosa* ATCC 27853 |
| Biofilm | This study | *P. aeruginosa* PA14 |

MIC=Minimum Inhibitory Concentration
OPKA=Opsonophagocytosis Killing Assay

**Figure S1.**
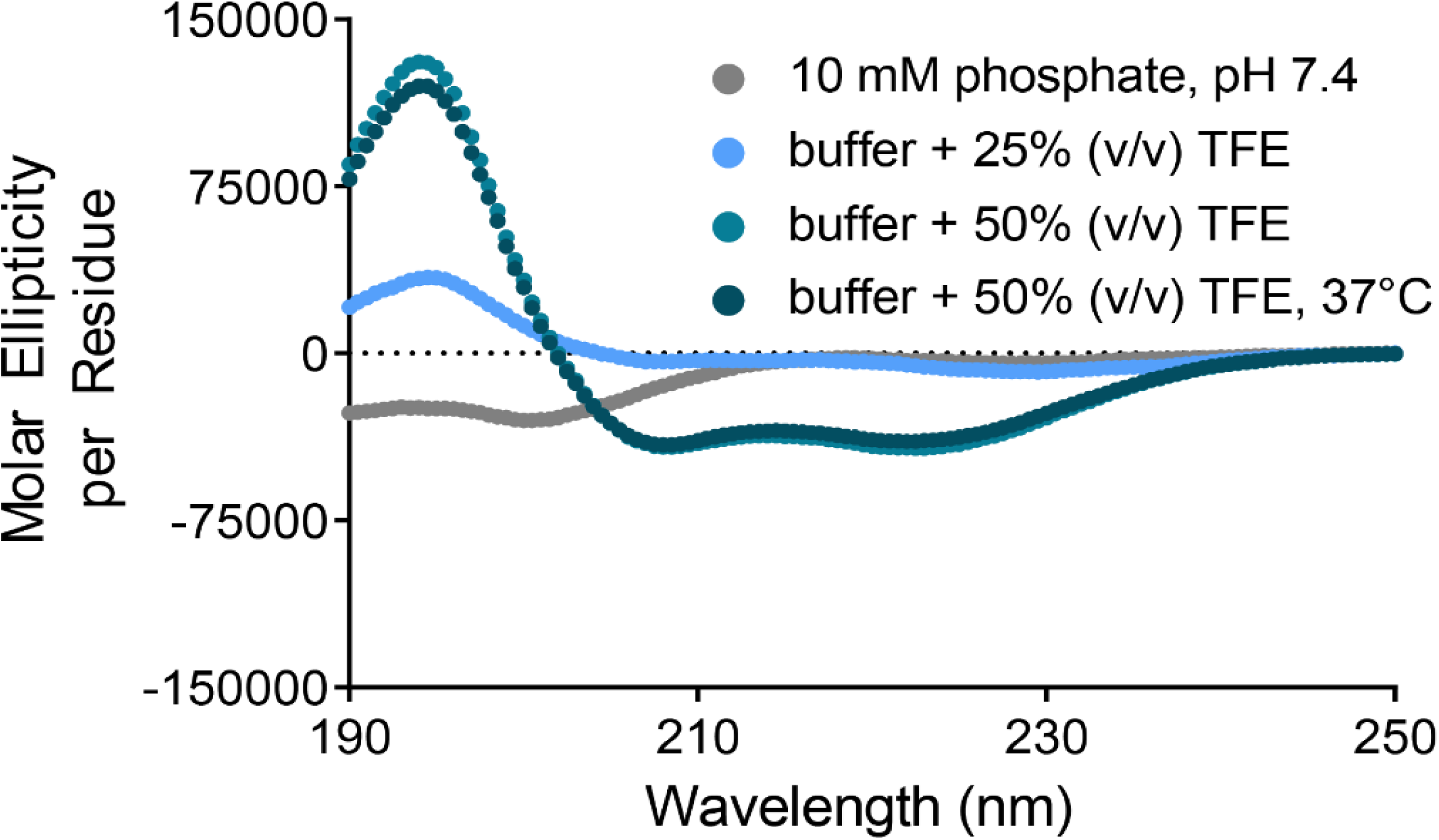
CD Analysis of P297. Circular dichroism spectra of P297 in buffer with increasing hydrophobic content. Graph shows increasing alpha-helical character with hydrophobicity.

**Figure S2.**
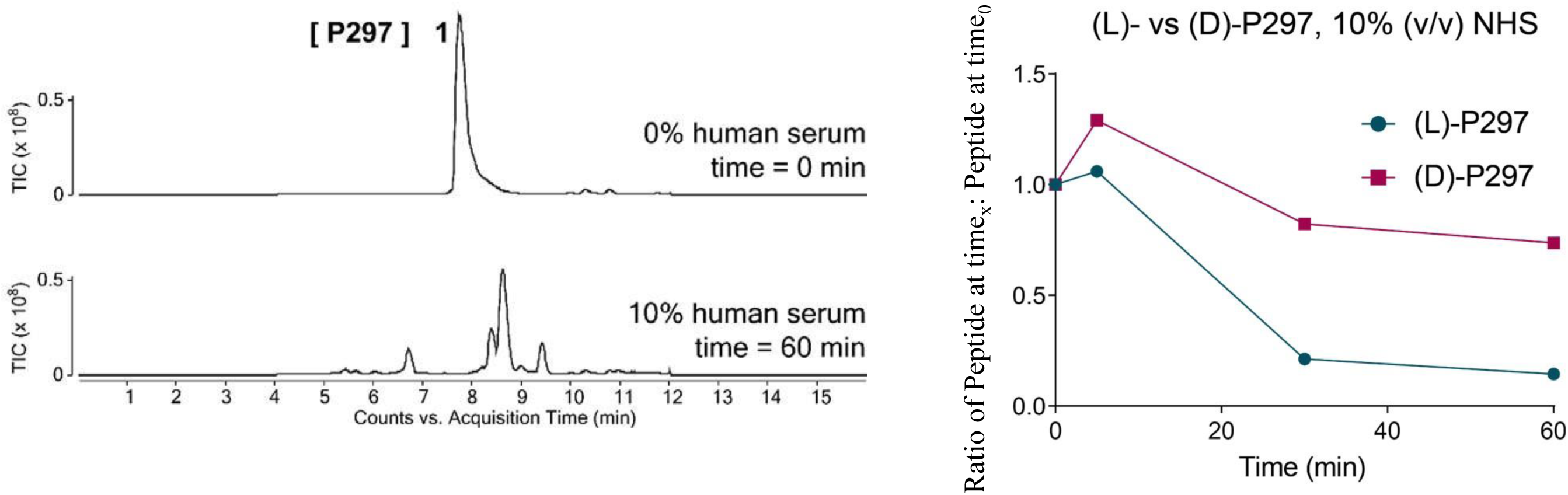
(D)-P297 is more stable than (L)-P297 in the presence of human serum. (left) Representative total ion chromatograms of P297 at t=0 (top) and after 60 minutes in 10% normal human serum (NHS). Intact peptide remaining was quantified by integration of extracted ion current for (L)- or (D)-P297. (right) Sixty-minute time course of degradation of P297 variants in the presence of 10% normal human serum (NHS). Values were normalized to the starting peptide content measured at time = 0 min.

**Figure S3.**
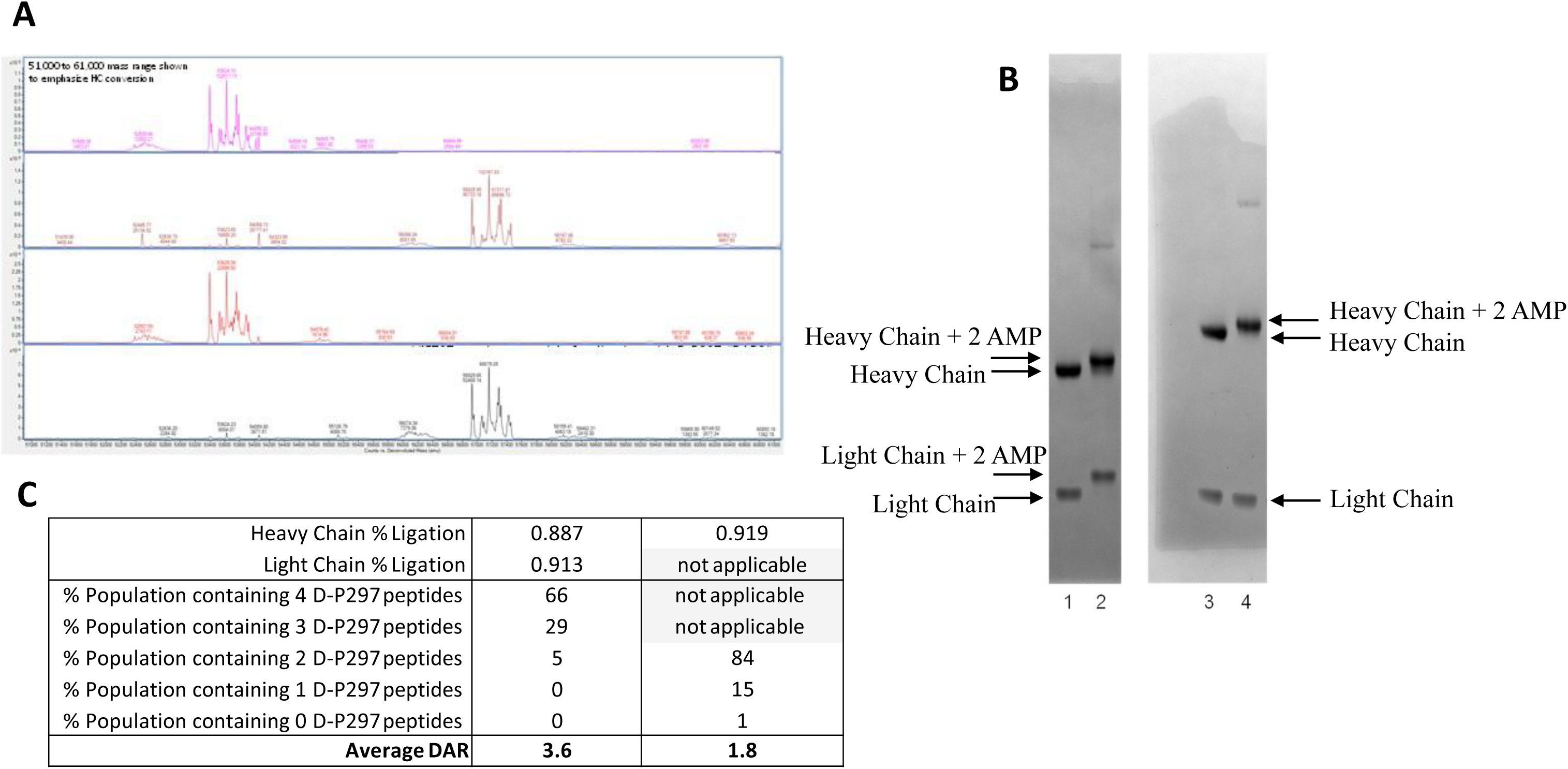
Analysis of Conversion of VSX to DAR2 and DAR4. (A) Q-tof analysis of the heavy chain of starting material and sortase-ligated D-P297 product for sortase acceptors on both the heavy and light chains (top two panels), and for sortase acceptors on only the heavy chains (bottom two panels). Assuming unbiased detection of starting material and product, 88.7% of the heavy chain for DAR4 material was ligated to D-P297. (B) Coomassie gel analysis showing conversion of starting material to desired D-P297 ADC product. Lanes 1 and 2 are the starting material and product for D-P297 ligation to both heavy and light chains. A second gel containing lanes 3 and 4 are respectively the starting material and product for D-P297 ligation to only the heavy chains (the light chain in the antibody starting material was wild-type lacking the sortase acceptor recognition sequence). (c) Calculation of the average DAR for constructs using deconvoluted spectra: with a theoretical DAR of 4, the experimentally determined DAR was 3.8; for the theoretical DAR of 2, the experimentally determined average DAR was 1.8.

**Figure S4.**
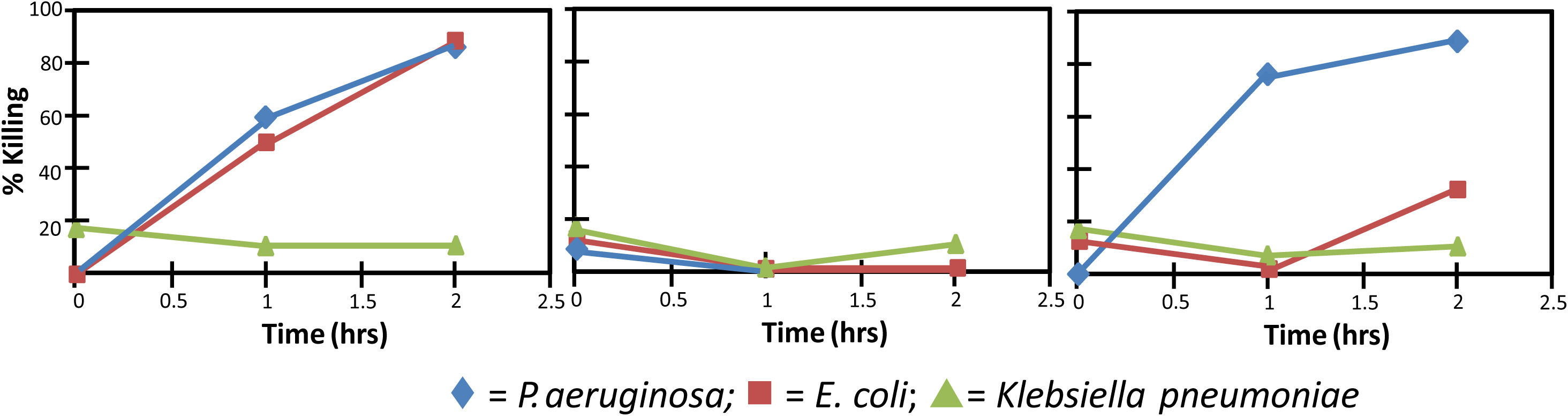
Mixed microbial killing assay. *P. aeruginosa* (blue), *E. coli* (red) and *K. pneumoniae* (green) were co-cultured overnight, diluted and then subjected to the specified agent for two hours. Killing was assessed visually. (left) Peptide P297 alone killing of *P. aeruginosa* and *E. coli*. (middle) In the absence of complement or immune cells, VSX alone had no killing effect. (right) VSX-1 demonstrated more rapid and complete killing of *P. aeruginosa*.

**Figure S5.**
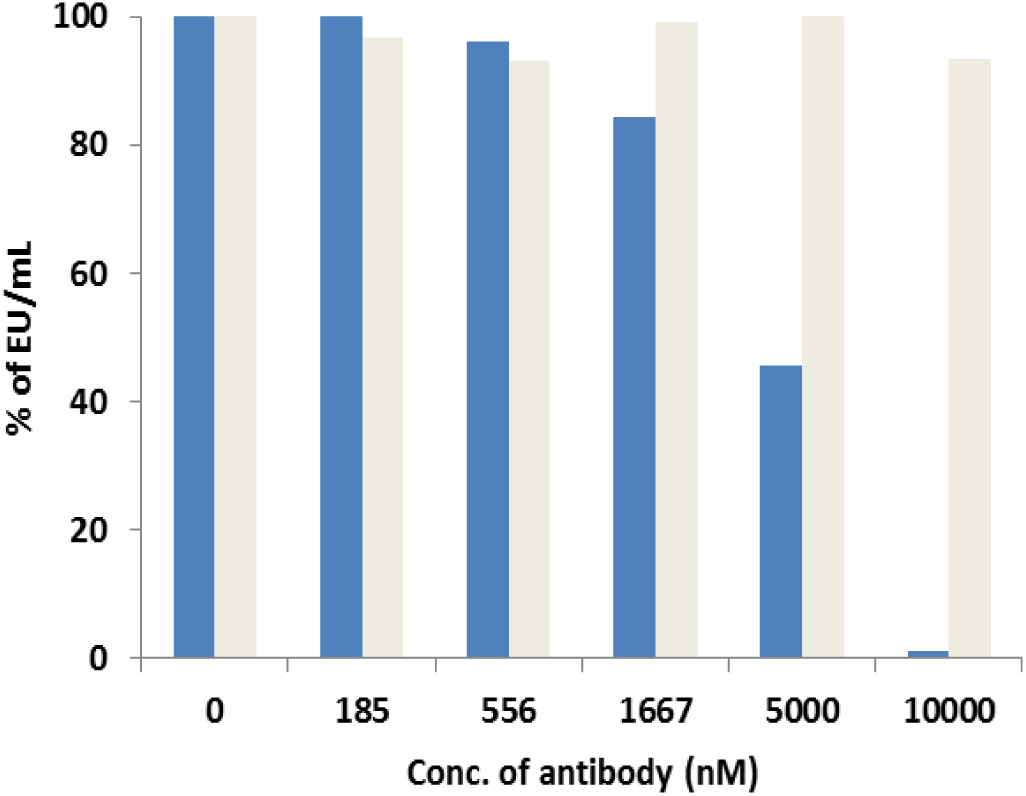
Neutralization of *P. aeruginosa* LPS activity in vitro. Binding of either VSX (blue) or actoxumab (grey) to LPS in a developed and optimized cell-based LPS neutralization assay. HEK-Blue LPS detection Kit was ordered from Invivogen to investigate the ability of VSX to neutralize endotoxin activity of extracted *P. aeruginosa* LPS on HEK-blue cells. In this case, endotoxin present in the media or standard is sensed by TLR4 leading to the activation of NF-kB and the production of SEAP in the supernatant. When supernatant is combined with QUANTI-Blue, this activation can be visualized and compared to a standard curve. 0.5 EU/ml *P. aeruginosa* LPS serotype was used for all assays.

